# Accurate plant pathogen effector protein classification *ab initio* with deepredeff, an ensemble of convolutional neural networks

**DOI:** 10.1101/2020.07.08.193250

**Authors:** Ruth Kristianingsih, Dan MacLean

## Abstract

Plant pathogens cause billions of dollars of crop loss every year and are a major threat to global food security. Effector proteins are the tools such pathogens use to infect the cell, predicting effectors de novo from sequence is difficult because of the heterogeneity of the sequences. We hypothesised that deep learning classifiers based on Convolutional Neural Networks would be able to identify effectors and deliver new insights. We built a training set of manually curated effector sequences from PHI-Base and used these to train a range of model architectures for classifying bacteria, fungal and oomycete sequences. The best performing classifiers had accuracies from 93 % to 84 %. The models were tested against popular effector detection software on our own test data and data provided with those models. We observed better performance from our models. Specifically our models showed greater accuracy and lower tendencies to call false positives on a secreted protein negative test set and a greater generalisability. We used GRAD-CAM activation map analysis to identify the sequences that activated our CNN-LSTM models and found short but distinct N-terminal regions in each taxon that was indicative of effector sequences. No motifs could be observed in these regions but an analysis of amino acid types indicated differing patterns of enrichment and depletion that varied between taxa. We have produced an R package that will allow others to make easy effector predictions using our models.

## Introduction

Phytopathogens are a major threat to global crop production. The fungal phytopathogen *Magnoporthe oryzae* that causes cereal blast is responsible for around 30% of rice production loss and has now emerged as a pandemic problem on wheat (Nalley et al., 2016) The oomycete *Phytophthora infestans* causes losses of around 6 billion USD to potato production, annually (Haas et al., 2009). The bacterium *Ralstonia solanacearum* has a wide host range and can cause loses of over 30% in potato, banana and groundnut (Yuliar, Nion and Toyota, 2015). The incidences of crop disease are increasing, global climate change and agricultural practice are expanding the geographical range of pathogens and upping the stakes in the evolutionary arms race. Effector proteins are the shock troops of infection, manipulating the host at the infection interface to the pathogens advantage. Identifying and characterising a pathogen’s effector content is a critical first step in understanding diseases and developing resistance, but effectors are notoriously difficult to characterise from sequence data. In most phyla they have only a few easily determined sequence characteristics (some in fungi are cysteine rich or have a MAX motif, some in oomycetes have the RXLR motif or WY fold) but in many cases no sequence identifiers are known (Franceschetti et al., 2017). Characterising effectors requires painstaking molecular experimental work and genome-scale approaches have relied on complex computational pipelines with in-built *a priori* assumptions about what might constitute an effector sequence in the absence of sequence features known to group them (Sperschneider et al., 2015). To understand infection processes, to provide genome-level understanding of the functions of this important class of genes and to develop future disease resisting crop varieties there is a need to identify effectors computationally from genome and protein sequence data.

Machine learning (ML) algorithms are a general group of techniques most often used for classification of data into groups. Supervised ML require a set of training examples and associated data with which to learn. Defining the best data to use and collect, called feature selection is an important and difficult prerequisite. ML approaches have been applied with success to biological sequence analysis, particularly in transcription factor binding site prediction, for the classification of eukaryote and bacterial nuclear proteins (Savojardo et al., 2017) and in the plant pathogen domain work by Sperschneider et al (Sperschneider, Dodds, Singh and Taylor, 2018) developed two ensemble-based machine learning models that could identify effectors and predict localisation with > 70% accuracy (Sperschneider et al., 2016, Sperschneider, Dodds, Gardiner, Singh and Taylor (2018)).

Deep learning models are distinct from other machine learning processes in that pre-selection of important features is far less critical and the models can learn these features unsupervised from training data (Jurtz et al., 2017). This property removes the need to know which properties of a data set must be examined before data collection begins. The Deep learning models can therefore classify on properties not necessarily known to the operator and could be used to uncover cryptic patterns in data. Convolutional neural networks (CNNs) are a type of neural network that have found wide application in numerous machine vision problems, including image object classification and facial identification (Krizhevsky, Sutskever and Hinton, 2012, Lawrence et al. (1997)), in time-series data analysis (Pyrkov et al., 2018) and natural language processing (Collobert and Weston, 2008). In the biomedical domain they have been used in drug discovery (Wallach, Dzamba and Heifets, 2015) and gene network prediction (MacLean, 2019). In studies with bacterial type III secreted effectors Xue et al developed an accurate CNN classifier for bacterial sequences (Xue et al., 2018). CNNs encode information about the features used to classify that can be extracted and interpreted. In a sequence classification problems this means they have the potential to reveal novel sequence features that other bioinformatics approaches have not and could be of particular utility when analysing sets of effectors.

Deep learning approaches require positive and negative examples from which to learn - here a list of sequences annotated as an effector or not. The larger and more accurate the list the more sensitivity a model can obtain. It is critical that training examples are experimentally verified effectors. Much of the effector annotation in public genomics databases is from computational predictions of genomics and is therefore of experimentally unverified hypothetical effectors. A good source of experimentally verified data is in the Molecular Plant Microbe Interactions (MPMI) literature and The widest ranging manual curation of MPMI papers is being performed as part of the PHI-Base (Urban et al., 2019) database strategy, PHI-base is an expertly curated database of genes proven experimentally to affect the outcome of pathogen host interactions and is therefore an excellent source of reliable effector sequences.

Here we use combinations of CNNs that do not rely on *a priori* feature selection to classify experimentally verified effectors and non-effectors in three taxa of plant pathogen: bacterial, fungal and oomycete. We show that these have very strong predictive power, can outperform existing effector prediction methods in accuracy and a better balance of sensitivity and specificity. We also analyse the activations of the models in response to effectors and non-effectors to gain insights into the sequence features that are allowing classification. We have produced an R package that will allow other scientists to easily classify their own sequences of interest.

## Methods

### Sequence Data Collection

Sequence data were collected from the PHI-Base database version 4.8 (Urban et al., 2019) by accessing a text dump of the data prepared on request by the PHI-Base team, the file can be accessed at https://github.com/PHI-base/data/blob/master/releases/phi-base_current.csv. The pipeline in Figure 1 outlines the steps used. We filtered plant effector proteins and their taxonomic groups and collected sequences from UniProt Release 2019_05, using the code in https://github.com/TeamMacLean/ruth-effectors-prediction/blob/master/scripts/r-scripts/getting-data-new/binary-class/0001_first_step_getting_data.Rmd. We created a correspondingly sized data set of non-effectors with secretion signals originating in species matched to those from which the effectors were drawn. We downloaded sequences for randomly selected proteins matching these criteria from Ensembl databases (Yates et al., 2020): specifically Ensembl Fungi, Protists and Bacteria manually using the BioMart tools (Smedley et al., 2009). Since the BioMart tool is not available on Ensembl Bacteria, we downloaded whole proteome protein sequnces from species matched to those from which the effector came using FTP. With these we used SignalP 3.0 (Dyrløv Bendtsen et al., 2004) in order to filter the secreted sequences and selected accordingly. We used default paramaters from SignalP, except the type of organism group which is euk for both fungi and oomycete sequences, and gram- or gram+ for bacteria sequences. Redundant sequences were filtered using BLASTp (Camacho et al., 2009). We achieved these steps using the code in https://github.com/TeamMacLean/ruth-effectors-prediction/blob/master/scripts/r-scripts/getting-secreted-data/0005_process_signalp_data.Rmd.

**Figure 1:**
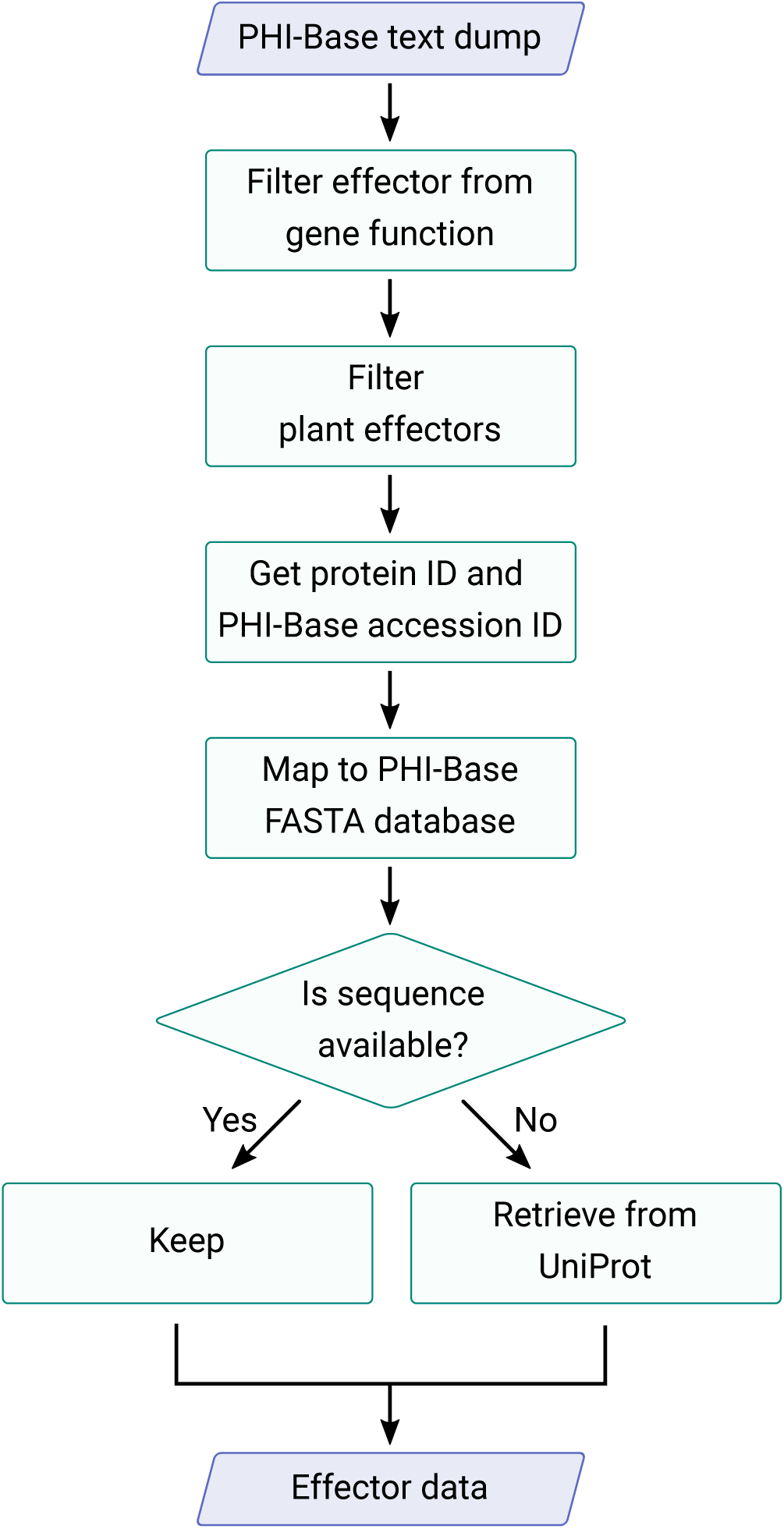
Workflow diagram for collection of effector sequences from the PHI-Base database annotation and cross reference to UniProt

### Encoding and Subsetting Sequences

The sequences collected were encoded using either one-hot encoding (CNN-LSTM based models) or integer based encoding (CNN-GRU-LSTM models). Sequences were post-padded with zeroes to bring the vectors to identical lengths to each other and the longest sequence in the taxon data set. The longest sequence for bacteria, fungi, and oomycete are 2574, 4034, and 934, respectively. Encoded sequences were split into taxon specific training, test and validation sets at a 60%, 20%, 20% split respectively as described in code at https://github.com/TeamMacLean/ruth-effectors-prediction/blob/master/scripts/r-scripts/getting-secreted-data/0008_split_and_encode.Rmd

### Model Training

We trained four model types on each taxon specific sequence set: CNN-LSTM, CNN-GRU, LSTM-Embedding, GRU-Embedding. We trained each model using a basic random hyperparameter setting initialisation step followed by hyperparamter scans. All models were implemented in Python 3.6.9 (Van Rossum and Drake Jr, 1995) using the deep learning API Keras 2.2.4 (Chollet et al., 2015) with the Tensorflow 1.12.0 (Abadi et al., 2015) backend, using NVIDIA GK110GL Tesla K20c GPUs and AMD Opteron(TM) Processor 6272 CPUs with 128 GB RAM.

### Hyperparameter Scans

Hyperparameter scans were performed using random search, a hyperparameter optimization method where each hyperparameter setting is randomly sampled from a distribution of possible hyperparameter values (Bergstra and Bengio, 2012). We used RandomSearchCV(), the implementation of random search in `scikit-learn 0.19.2` (Pedregosa et al., 2011) together with KerasClassifier() which is an implementation of the scikit-learn classifier API for keras. Code for this can be found in https://github.com/TeamMacLean/ruth-effectors-prediction/tree/master/scripts/python-scripts/hyperparameter-scan-scripts. All model training was performed as described in section Model Training.

### Fine tuning

Fine tuning was performed manually using `keras 2.2.4` together with metrics module and KFold cross validator from `scikit-learn 0.19.2`. Scripts implementing the tuning can be found at https://github.com/TeamMacLean/ruth-effectors-prediction/tree/master/scripts/python-scripts/manual_tune_scripts.

### Model Classification Correlation

We calculated correlations between the classifications from best performing models on the hold-out test data set using Pearson’s correlation co-efficient on the 1/0 classification vectors.

### Ensemble Functions

We computed an aggregate classification using two different ensemble functions, weighted average and an overall majority option.

Weighted average is computed as

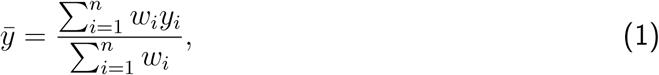

where *w*_*i*_ is the weight, *y*_*i*_ is the prediction value of the *i*^*th*^ model, and *n* is the total number of model. In our case, we use the accuracy of each model as the average.

Overall majority is computed as

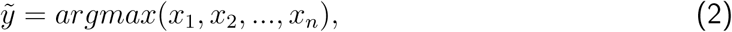

where

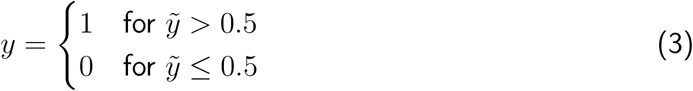

### Metrics

We used the following calculations for different accuracy metrics in our evaluations, specifically: accuracy, sensitivity, specificity. *TP, TN, FP*, and *FN* refer to the number of true positives, true negatives, false positives and false negatives, respectively.

Accuracy (*Acc*) is the ratio between correctly classified non-effectors and effectors and all samples:

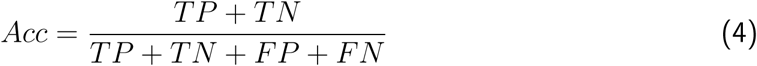

Sensitivity *Sn* is the ratio between correctly predicted as effectors and all effectors:

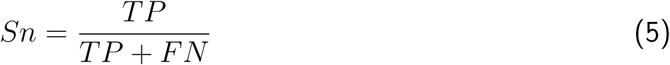

Specificity (*Sp*) is the ratio between correctly predicted as non-effectors and all non-effectors:

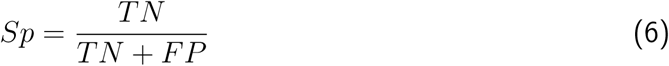

F1-score is the harmonic average of the precision and recall:

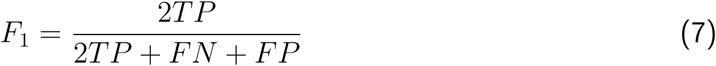

### Activation Map Analysis

To visualise the regions of the sequences that the models were using to discriminate between effector and non-effector we adapted a Grad-CAM (Gradient-weighted Class Activation Mapping) approach (Selvaraju et al., 2019). We extracted the convolutional layer convd_1 and computed the gradient of the model output with respect to the output of convd_-1. The feature map was weighted by every channel and mean result taken. All activation maps were summed, normalised and smoothed using Fourier analysis. We used tensorflow 1.12.0 to compute the activation maps and discrete fourier transform (numpy.fft) from numpy 1.17.3 to smooth the result. The code we used for computing and visualising these heatmaps can be found at https://github.com/TeamMacLean/ruth-effectors-prediction/tree/master/scripts/python-scripts/heatmaps.

### Effector Prediction Software and Training Data

To test the performance of our models against commonly used tools we used DeepT3 version 1 (Xue et al., 2018) and EffectiveT3 version 1.0.1 (Eichinger et al., 2015) for bacterial sequences. Effector P Version 1.0 and 2.0 (Sperschneider et al., 2016, Sperschneider, Dodds, Gardiner, Singh and Taylor (2018)) for fungal sequences and EffectR (Tabima and Grünwald, 2019) for oomycete sequences. All the models publish the datasets used to train the models or some examples. We used positive training examples (effector sequence examples) in the comparisons we performed. EffectorP provides three different positive datasets (training, test and hold-out validation) for EffectorP 1.0 and 2.0 at http://effectorp.csiro.au./data.html. EffectiveT3 provides a training set at https://effectors.csb.univie.ac.at/sites/eff/files/others/TTSS_positive_training.faa. DeepT3 provides three sets, a non redundant Pseudomonas syringae effector dataset, a training dataset and a test data set at https://github.com/lje00006/DeepT3/tree/master/DeepT3/DeepT3-Keras/data. EffectR uses 6 RXLR oomycete sequences described in (Haas et al., 2009) as examples rather than as a training set, namely *PexRD36, PexRD1, ipi01/Avrblb1, Avr1, Avr4*, and *Avr3a*. All these sets were used in their respective tools with default settings.

## Results

### Sequence Collection

The performance of the trained classifiers is dependent on the quality of the input training data, so it was important that we collected as high a quality set of annotated effectors as possible.

To this end we used PHI-Base (Urban et al., 2019) as our primary sequence origin. Sequences in PHI-Base are human curated from the literature and have therefore been noted in experimental studies. They do not derive from large scale annotations or contain hypothetical or predicted proteins. This attribute makes it ideal for our purposes as the effectors in PHI-Base are those that have been specifically reported as such in the published literature and are not of the class of sequences that are merely suspected of being effectors on the basis of carrying a secretion signal. To collect effector sequences we parsed a whole database text dump of version 4.8 https://github.com/PHI-base/data, all proteins marked as plant pathogen effectors were filtered and we used the IDs and UniProt IDs to collect the protein sequences from PHI-Base or UniProt if PHI-Base stored only the ID. The sequences and IDs retrieved can be seen in the data file in this manuscript’s repository https://github.com/TeamMacLean/ruth-effectors-prediction/blob/master/data/getting-data-new/binary-class-data/effector_data.csv. Effector sequences were then divided into taxonomic groups as bacterial, fungal or oomycete derived accordingly. In total 190 bacterial effectors from 13 species were collected, 97 fungal effectors from 16 species were collected and 85 oomycete effectors from 6 species were collected (Table 1). The species and effector count in each group can be seen in Tables S1, S2 and S3.

**Table 1:**
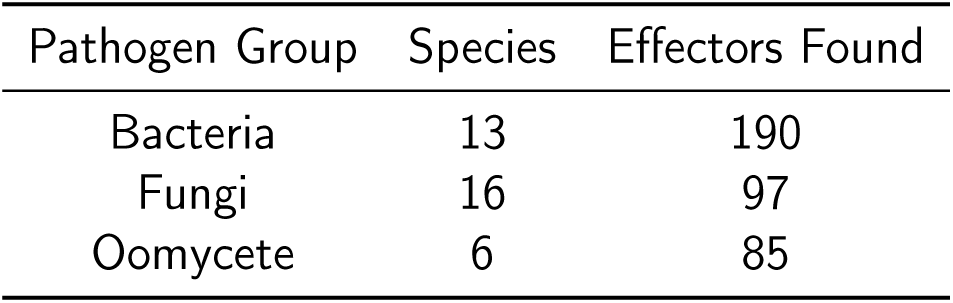
Count of effectors listed in publications curated by PHI-Base used in this study in three major plant pathogen groups

Sequences for non-effector, secreted proteins were collected using a similar pipeline. Randomly selected proteins from each species carrying secretion signals were extracted from Ensembl databases using the BioMart tool. For each species noted in Tables S1, S2 and S3 we collected from either the same strain or species an identical number of non-effector, secreted proteins to that in the effector set. This gave us a balanced data set of effector proteins as positive learning examples and non-effector secreted proteins as negative learning examples. Figure 2 summarises the process of building the non-effector set, and the full set of sequences and IDs retrieved can be seen in the following data file https://github.com/TeamMacLean/ruth-effectors-prediction/tree/master/data/secreted_data.

**Figure 2:**
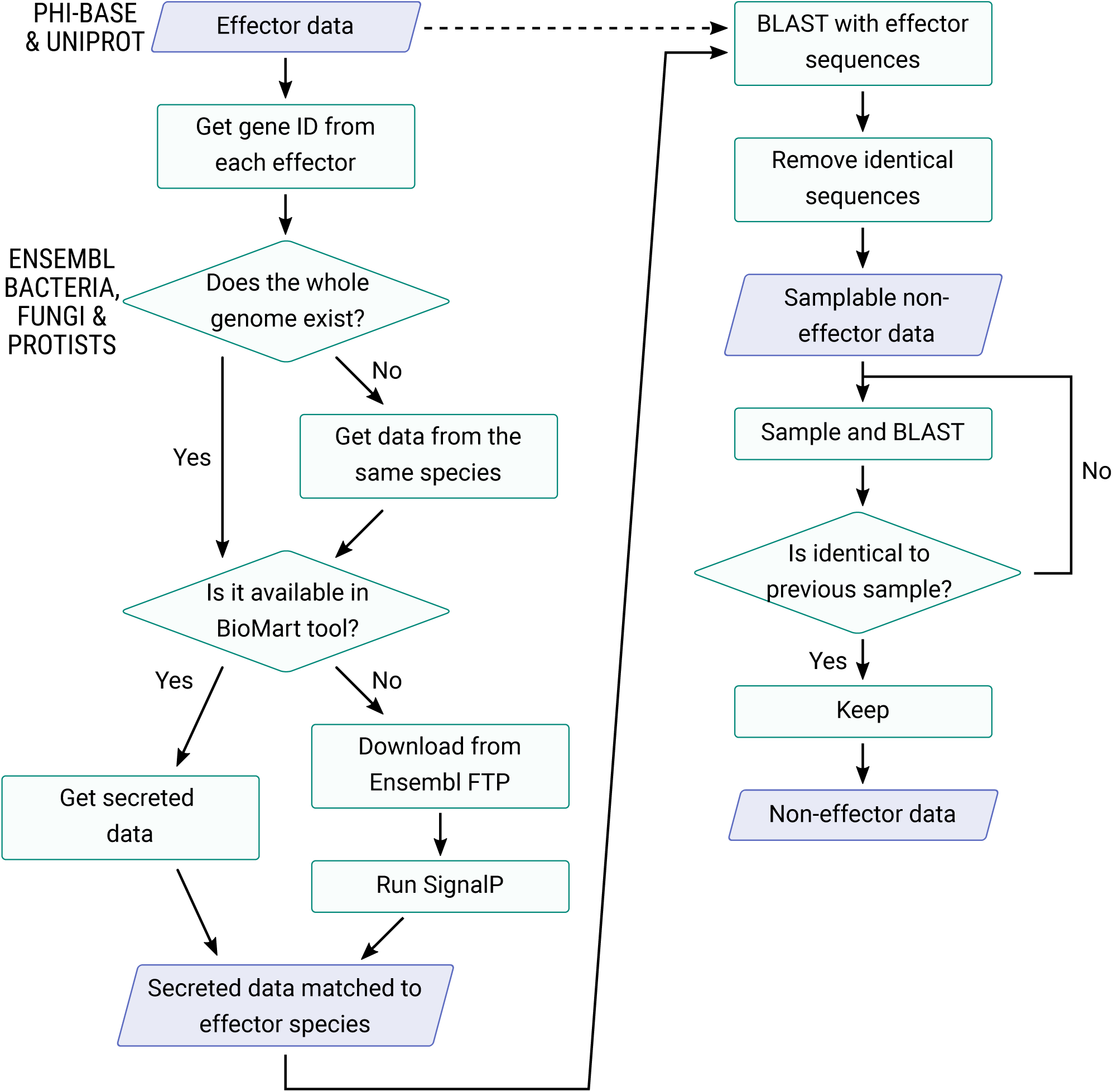
Workflow diagram for collection of secreted non-effector sequences from Ensembl Bacteria, Fungi, and Protists

### Model Selection and Training

In order to identify a useful classifier we took a randomised hyperparamter search over some likely base model architectures. We selected four base architectures on which to build models for learning. Two of these contained Convolutional Neural Network (CNN) layers followed by either a Long Short Term Memory Layer (LSTM) or a Gated Recurrent Unit (GRU), two contained an Embedding Layer followed by the LSTM or GRU. All models had fully-connected dense layers after this. See Figure 3.

**Figure 3:**
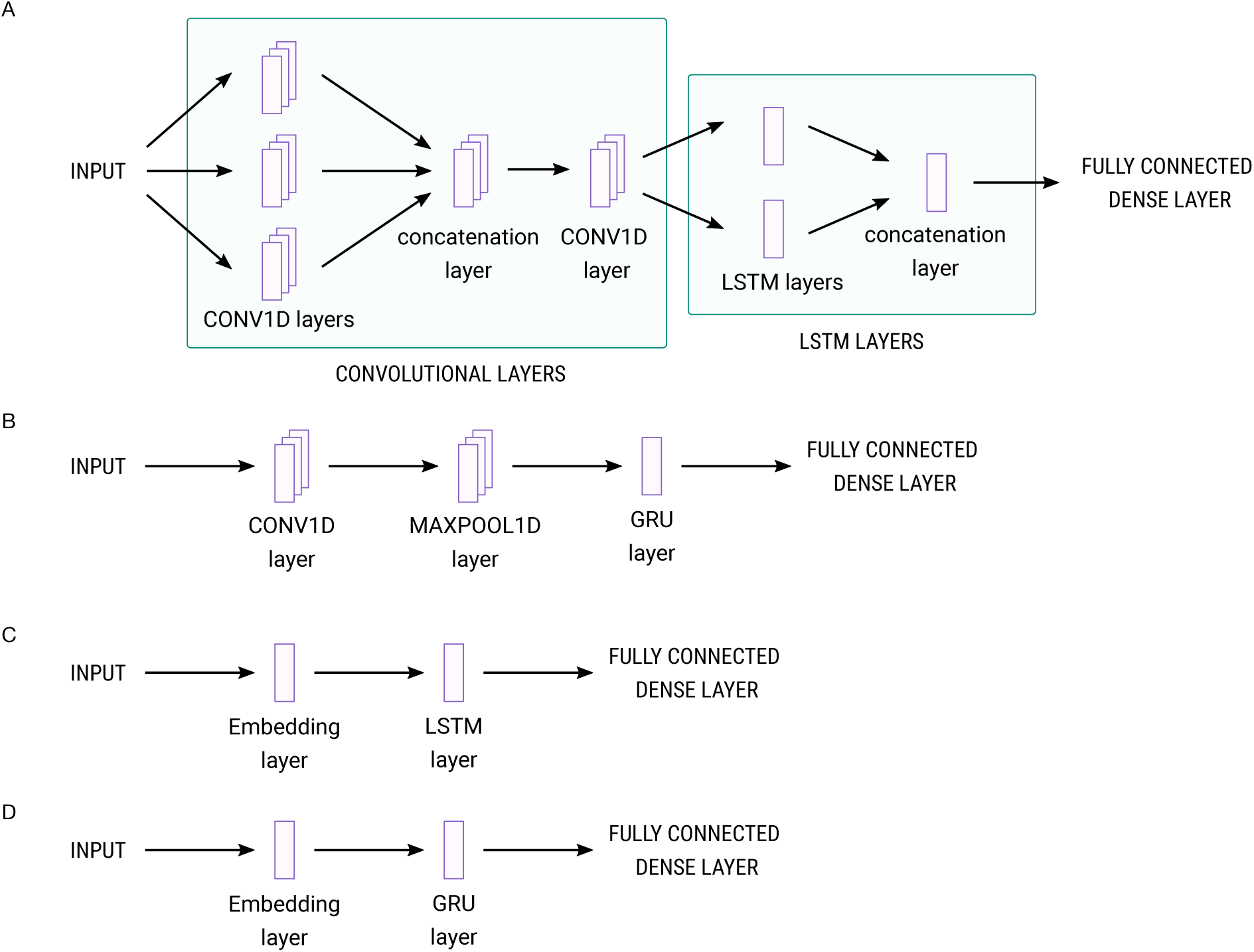
Overview of model architectures tested in this analysis, A: CNN-LSTM model; B: CNN-GRU model; C: LSTM-Embedding model; D: GRU-Embedding model

We defined a range of values for the hyperparameters that could be optimised in each architecture, 10 for CNN-LSTM, 12 for CNN-GRU, 9 for LSTM-Embedding and 9 for GRU-Embedding. To test all combinations of values in these ranges would take a prohibitive amount of processor time, so we selected 50 sets of values for each model in each taxon at random to start training, 3000 models in total. Model variants within the hyperparameter search were assessed by comparing accuracy values on the development validation fraction of the training data. Other hyperparameters were fixed and are listed in Table S5. For each model type and taxon training data combination we selected the hyperparameter set giving highest accuracy on the validation fraction. From this we had twelve candidate models to develop further.

We then manually ran and checked the accuracy and loss of the twelve models on the training and validation sets to investigate instances of overfitting and assess generality. Smaller models are less likely to overfit data, so we investigated the effect of regularization rate, filter count and kernel size on the progress and accuracy of the model as we reduced the size. Parameters varied in this phase are listed in S6. Final selected hyperparameter settings for models in each taxon can be seen in Table S7. The values of accuracy and loss of each model produced are shown in Table 2. We found that by reducing the number of kernels on all models from 2 to 1 and the number of filters reduced from 32 to 16 we removed apparent overfitting and retained high accuracy, with training completing in 40 epochs.

**Table 2:** Accuracy and loss values from best performing parameters values for each model

| Group | Data set | Model | Accuracy | Loss |
| --- | --- | --- | --- | --- |
| Bacteria | Training | CNN-LSTM | 0.917 | 0.334 |
| Bacteria | Training | CNN-GRU | 0.965 | 0.148 |
| Bacteria | Training | LSTM-Emb | 0.978 | 0.129 |
| Bacteria | Training | GRU-Emb | 0.947 | 0.202 |
| Bacteria | Validation | CNN-LSTM | 0.855 | 0.377 |
| Bacteria | Validation | CNN-GRU | 0.947 | 0.205 |
| Bacteria | Validation | LSTM-Emb | 0.934 | 0.232 |
| Bacteria | Validation | GRU-Emb | 0.895 | 0.385 |
| Fungi | Training | CNN-LSTM | 0.907 | 0.362 |
| Fungi | Training | CNN-GRU | 0.814 | 0.434 |
| Fungi | Training | LSTM-Emb | 0.771 | 0.505 |
| Fungi | Training | GRU-Emb | 0.788 | 0.484 |
| Fungi | Validation | CNN-LSTM | 0.526 | 0.923 |
| Fungi | Validation | CNN-GRU | 0.579 | 0.799 |
| Fungi | Validation | LSTM-Emb | 0.711 | 0.504 |
| Fungi | Validation | GRU-Emb | 0.842 | 0.468 |
| Oomycete | Training | CNN-LSTM | 0.657 | 0.626 |
| Oomycete | Training | CNN-GRU | 0.833 | 0.429 |
| Oomycete | Training | LSTM-Emb | 0.863 | 0.314 |
| Oomycete | Training | GRU-Emb | 0.706 | 1.487 |
| Oomycete | Validation | CNN-LSTM | 0.735 | 0.761 |
| Oomycete | Validation | CNN-GRU | 0.618 | 0.677 |
| Oomycete | Validation | LSTM-Emb | 0.588 | 0.910 |
| Oomycete | Validation | GRU-Emb | 0.500 | 1.486 |

Final training progressions for each model in each taxon can be seen in Figure 4. We tested the finalised models on the hold-out test fraction of the data that had not been previously seen, for the four bacterial sequence trained models we had accuracies in the range 93.4 to 97.4, for the four fungal models we observed accuracy in the range 60.5 to 84.2 and for the four oomycete models we observed accuracy from 64.7 to 82.3, reported in Fig 5. All the models we generated had high statistical power and can accurately and reliably classify effectors from other secreted proteins in that taxon.

**Figure 4:**
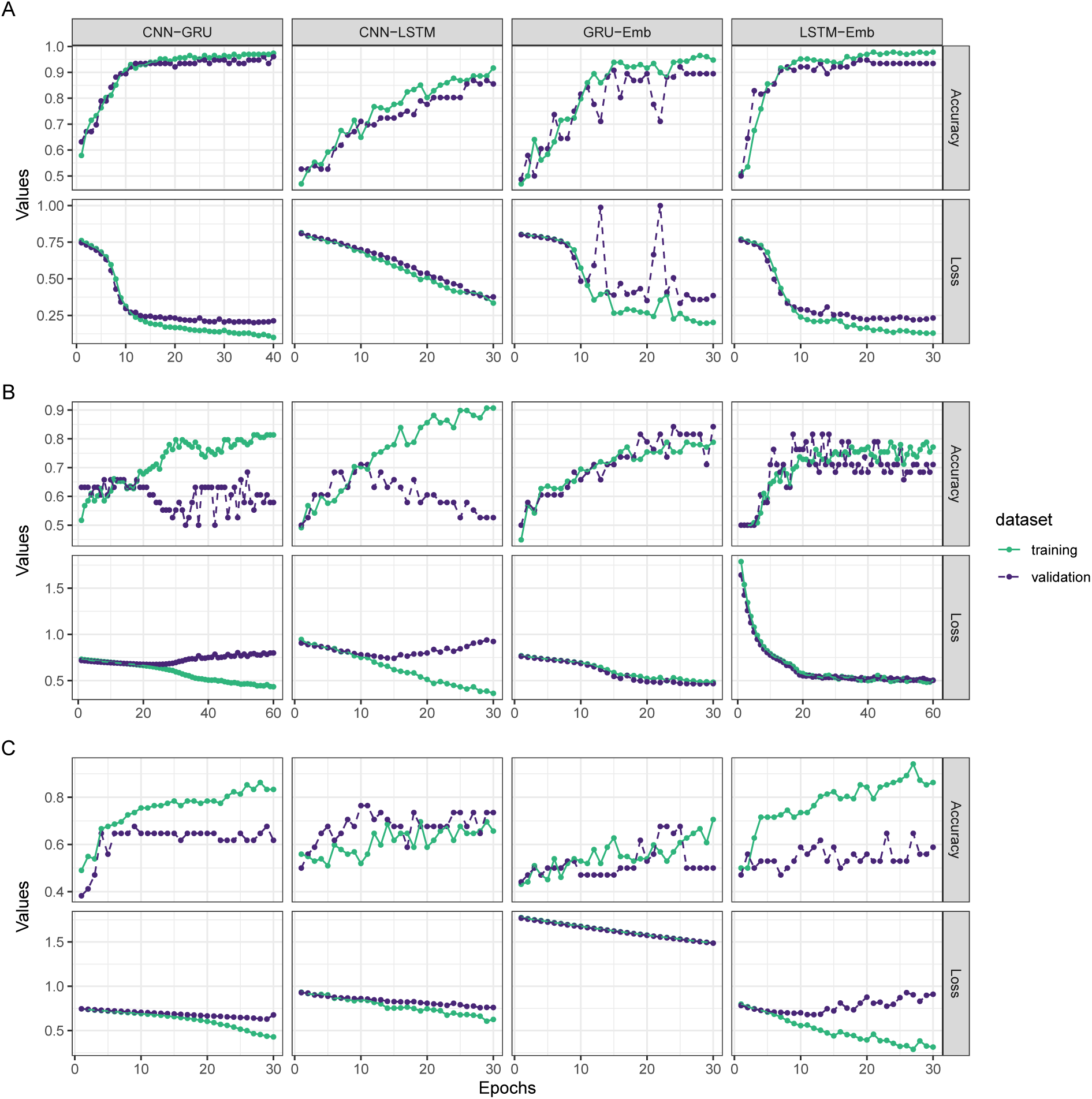
Training trajectories showing Accuracy and Loss over 40 epochs for four model types on A: Bacterial sequence training set; B: Fungal sequence training set; C: Oomycete sequence training set

**Figure 5:**
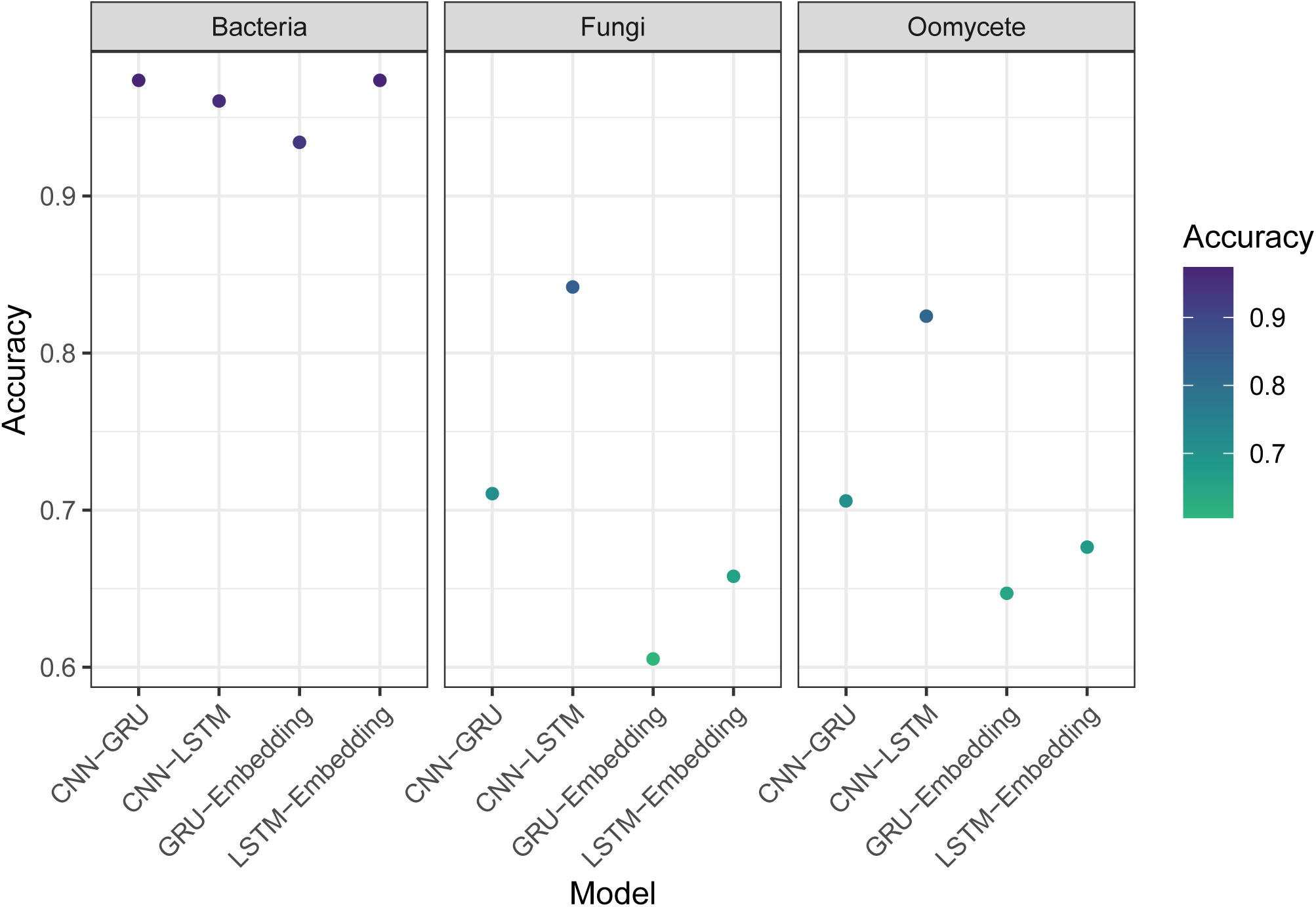
Final performance of models in each taxon on the unseen, hold-out test data fraction of sequences.

The final twelve models were saved into HDF5 objects and stored in the repository at https://github.com/TeamMacLean/ruth-effectors-prediction/tree/master/data/final_model_hdf5.

### Model Characteristics

We examined the tendency of the models to call false positives or false negatives preferentially by creating confusion matrices of the classifications relative to the ground truth on the hold-out test data. The bacterial sequence trained models in general showed high accuracy and only one or two miscalls with no error bias except for the GRU-Embedding model which called five from 38 effectors as non effectors. The fungal sequence trained models were less accurate overall and showed a small amount more bias, again in the GRU-Embedding model, which was biased towards calling effectors as non-effectors and the CNN-LSTM model which was slightly biased in the opposite direction, calling non-effectors as effectors. The oomycete models were again quite balanced but the GRU-Embedding model showed a quite conservative tendancy calling 12 out of 17 effectors as non-effectors whilst getting all 17 non-effectors correct. Overall the models are accurate and show little to no bias toward false positive or false negatives, with the exception of the GRU-Embedding type. In oomycete sequences in particular and in this class of model across the different sequence types showed itself to tend to call real effectors as not of that class.

Classification correlations between the different model architectures were high and positive in the bacterial sequence trained model’s calls, in the range 0.8 to 0.88, see Figure 6. CNN-GRU and LSTM-Embedding showed identical prediction sets. We observed similar levels of correlation in the CNN-LSTM, GRU-Embedding and CNN-GRU fungal sequence trained model, in the range 0.79 to 0.9; though there was a significantly lower range of correlations with the LSTM-Embedding which were in the range 0.36 to 0.51. The models trained on oomycete sequences all showed this lower range of correlations, in the range 0.30 to 0.65. The higher correlation across bacterial trained models is likely from a mixture of the larger training set size and a greater uniformity of the sequences themselves. For the fungal sequence trained models we can see that the LSTM-Embedding model does not perform as well as the others. The oomycete sequence trained models all show a lower range correlation reflecting the likely less uniform and smaller training set. It is clear that, particularly for the fungal and oomycete models, each architecture is capturing separate aspects of the sequences and classifying on those with slightly varying levels of success.

**Figure 6:**
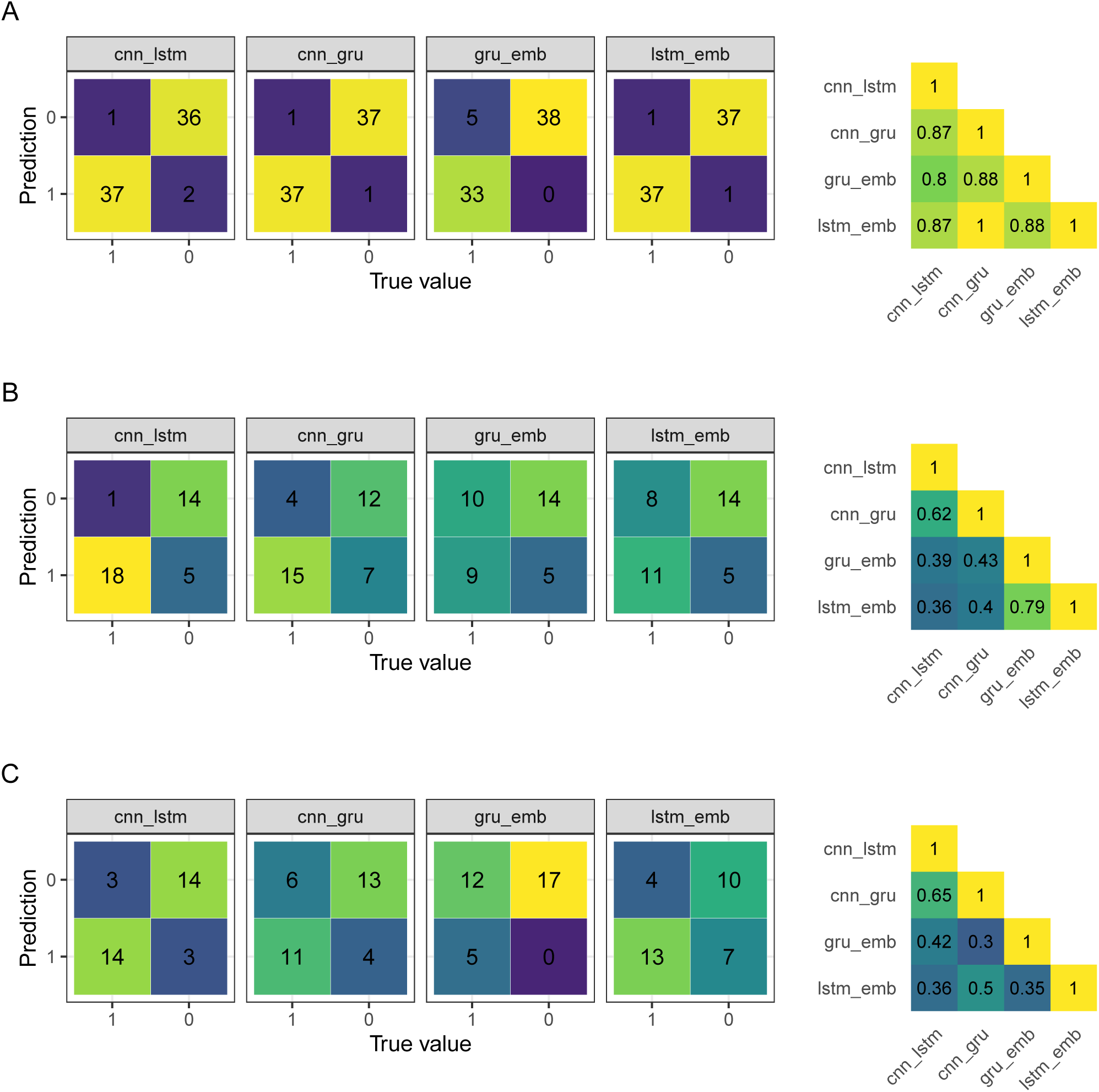
Confusion matrices showing tendency of models to call false positive or false negative errors on the final test data; and Pearson correlations between classifications made by different models. A: Models trained on bacterial sequences; B: Models trained on fungal sequences; C: Models trained on oomycete sequences.

### Ensemble Models

We examined the usefulness of combining the predictions of the different model architectures using an ensemble function that takes the vectors of classifications of each model architecture as input. We performed the classification of the hold-out test data set using the ensembled models and the results can be seen in Figure 7. With the models trained in bacterial sequences we observed an increase in classification accuracy over the best model, up to 0.99 for both ensemble functions. However, with the fungal and oomycete models we observed decreases relative to the best single model in both cases due to the higher accuracy of the CNN-LSTM model being diluted by the combined inacuracy of the other model architectures. Examining the overlaps in classifications between the CNN-LSTM and CNN-GRU/LSTM-Embedding respectively showed that the two lesser performing models were not simply predicting subsets of the CNN-LSTM model, in both cases the lesser models were able to identify three effectors correctly that were missed by the generally stronger models. This indicates that the weaker models may be classifying on some patterns missed by the CNN-LSTM model.

**Figure 7:**
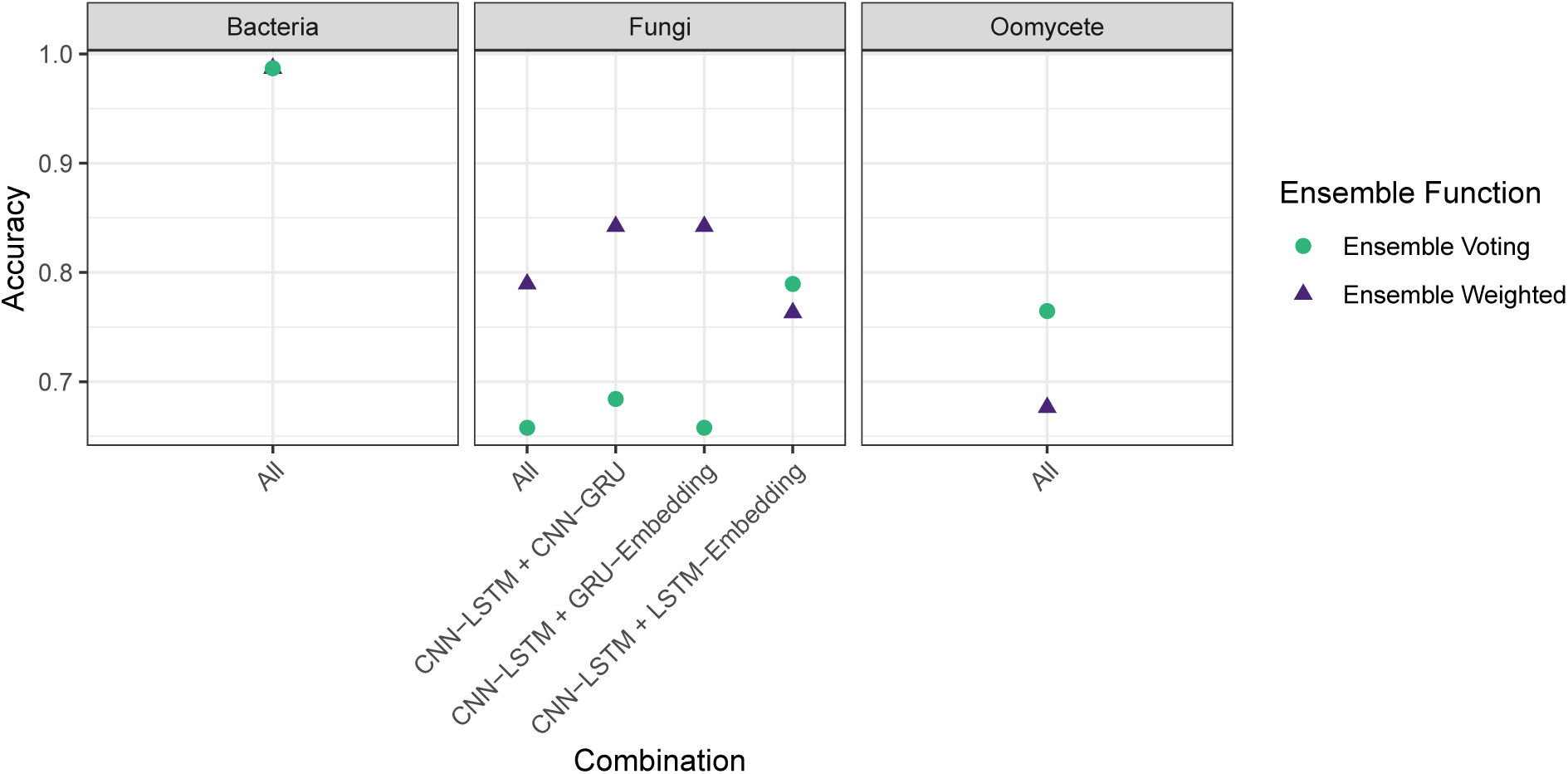
Final performance of Ensemble models in each taxon on the unseen, hold-out test data fraction of sequences.

### Comparison With Other Classification Software

Given the accuracies of the above we selected the ensemble bacterial model and the CNN-LSTM fungal and oomycete models to evaluate the performance of our models against widely used effector identification software. We compared against predictions from the bacterial effector prediction programs DeepT3 (Xue et al., 2018) and EffectiveT3 classification module for plant-associated bacteria 1.0.1 (Eichinger et al., 2015), the fungal effector prediction programs EffectorP 1.0 and 2.0 (Sperschneider et al., 2016, Sperschneider, Dodds, Gardiner, Singh and Taylor (2018)) and the oomycete effector predictor EffectR (Tabima and Grünwald, 2019). Each comparison was carried out using the respective hold-out test sequence set for each taxon. For all taxa we observed greater Accuracy and F1 scores from our models than the established software, as shown in Figure 8. This was particularly marked in the F1 score, which incorporates a measure of the incorrect calls. Absolute improvements were up to 15 % in bacterial sequences, 15 % in fungal sequences and 20 % in the oomycete sequences. The confusion matrices in Figure 9 show that accuracy and F1 score was compromised in all the established tools by the tendency of them all to misclassify true effectors as not effectors. All the established software classifiers we tested show lower sensitivity than the models we have developed here.

**Figure 8:**
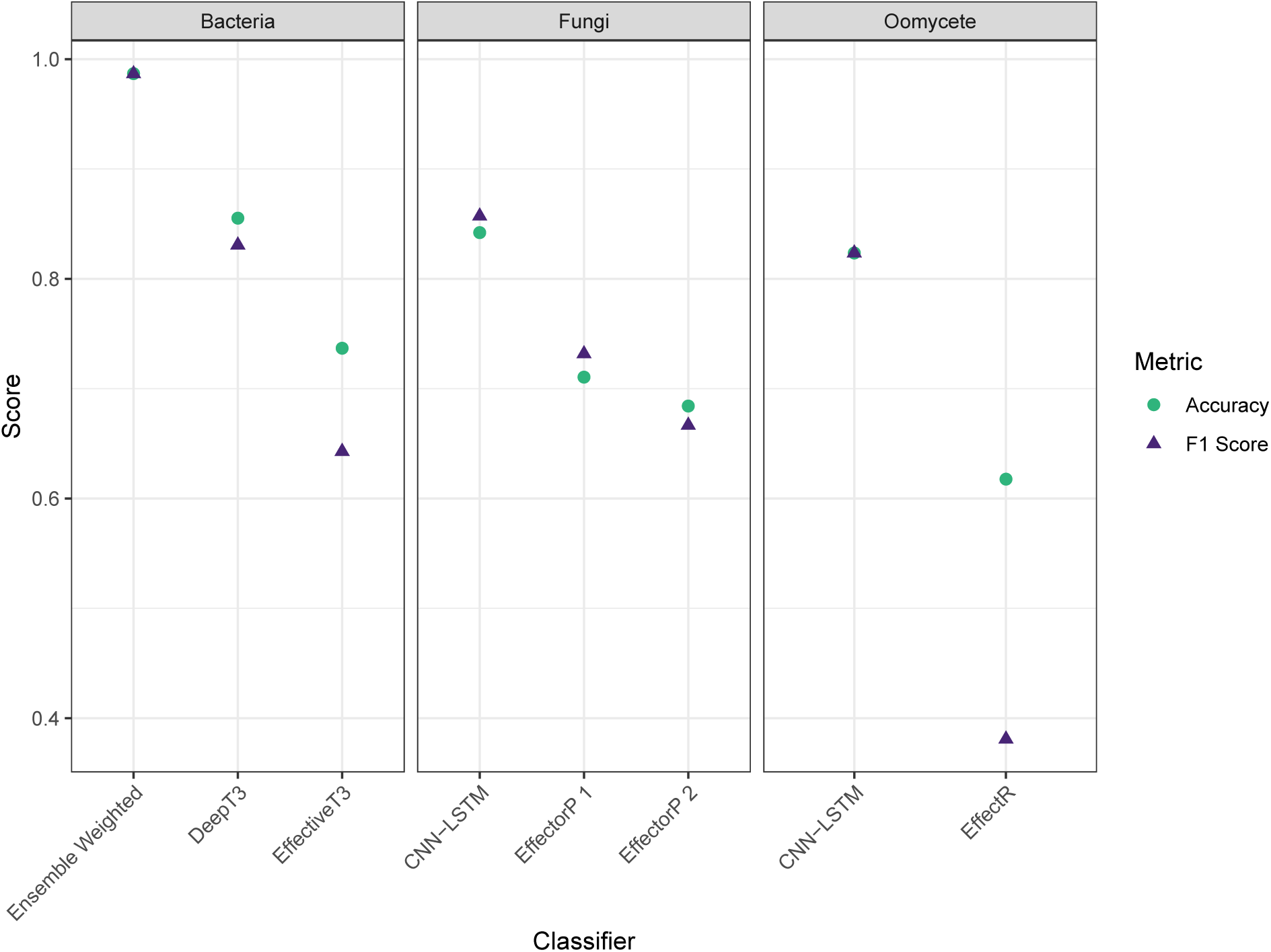
Accuracy and F1 score of the classifications made by our models compared with those made by widely used software

**Figure 9:**
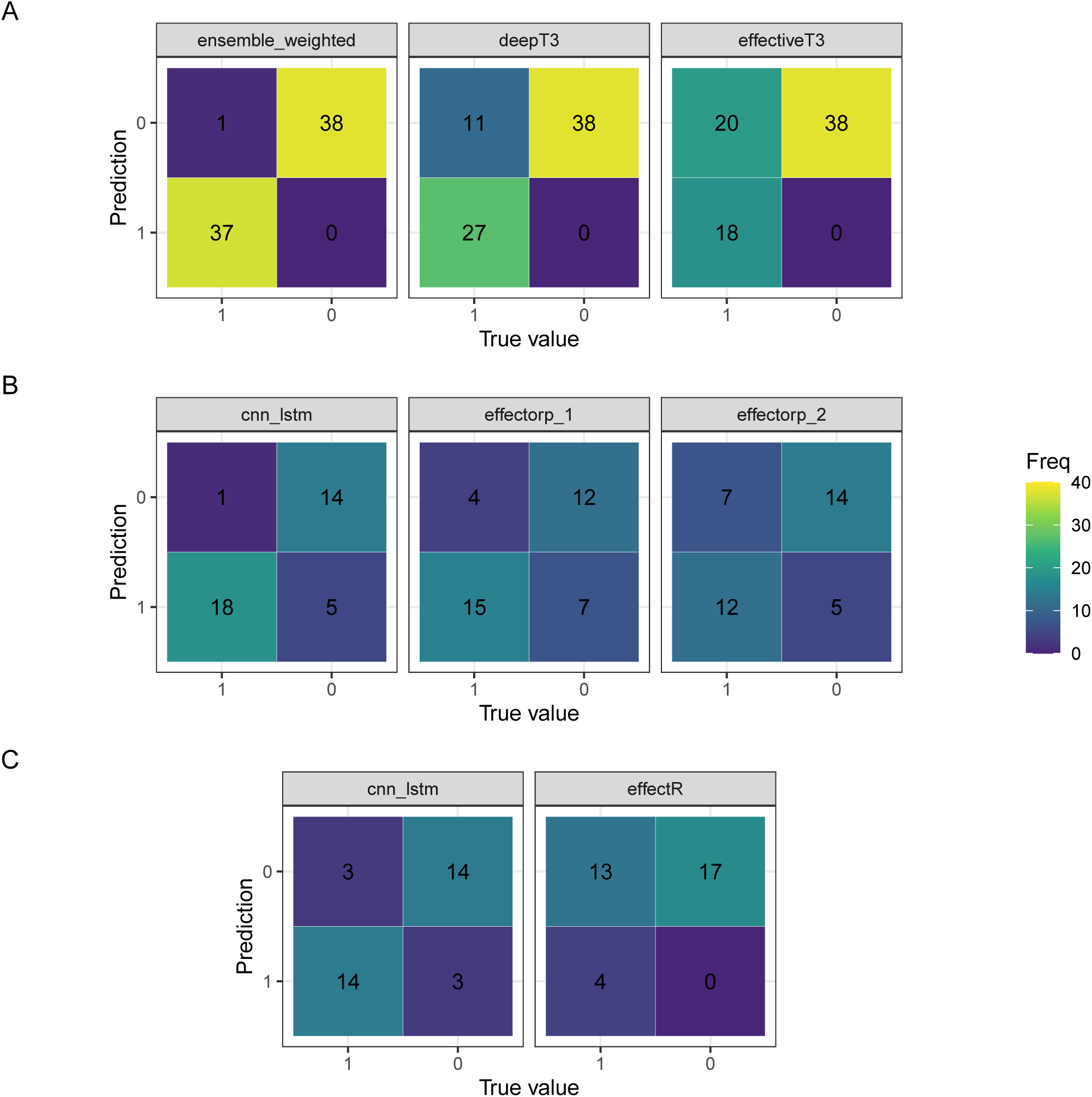
Confusion matrices of classifications made by our models compared with those made by widely used software. A: Tools evaluated on bacterial sequences; B: Tools evaluated on fungal sequences; C: Tools evaluated on oomycete sequences.

We also evaluated the deep learning models we have developed on the training and hold-out validation sequences used to train the previous methods. We calculated the proportion of the effectors in the training set that the tools could find on their respective training and validation sets, according to availability. The bacterial tools EffectiveT3 and DeepT3 showed lower proportion found than our Ensemble Weighted model, as in Figure 10A, consistent with the observation that our Ensemble Weighted model performed more strongly on the validation set that we generated. Interestingly, both versions of EffectorP found a greater proportion of the effectors in the EffectorP provided training sets than our CNN-LSTM model, but in the unseen validation data provided with EffectorP 2, all three models performed identically (Figure 10B). The EffectorP 1 and 2 scores on validation data are well below the scores for the training data, a result that is usually interpreted as being evidence of an overfitted and less generalisable model. Our CNN-LSTM model for fungi showed similar scores across the training and validation set indicating a greater generability and equivalent power. Only six effector sequences are provided as examples with the oomycete specific effector finder effectR. As this is not a trainable model in the same sense as the others, no large training set is needed. We attempted classification with these and effectR was able to classify 5 of the 6, whereas our CNN-LSTM model for oomycetes classified 3 of the 6 (Figure 10C).

**Figure 10:**
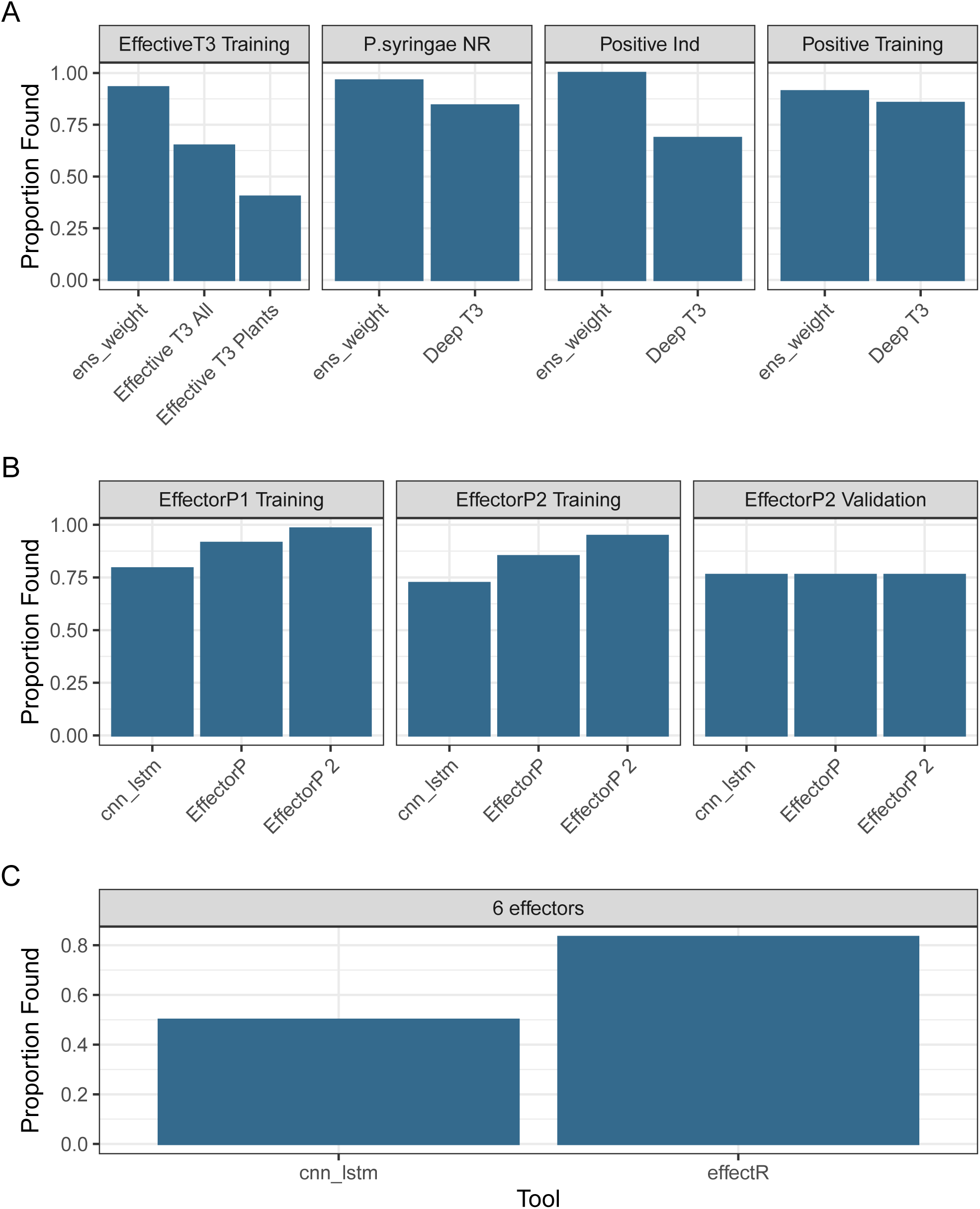
Performance of models on training and validation datasets used to train other effector prediction tools. A: Tools trained on bacterial sequences; B: Tools trained on fungal sequences; C: Tools trained on oomycete sequences.

Overall, our models performed more strongly than the previously available ones tested across the range of sequences examined.

### Convolution Heatmaps

An advantage of CNNs relative to other deep neural networks is their relative interpretability. A CNN can be analysed and activation maps extracted which highlight the regions of the input to which the CNN is most strongly relying on to classify. To examine the responses of the models, we ran all sequence data sets back through the CNN-LSTM models for each taxon and extracted the network activations using the GRAD-CAM method. The profiles were smoothed using FFT and examined, (see Figure 11). All the models showed a peak of activation at the N-terminus of the sequences, coincident with expected positions of secretion signals. The fungal sequences created a single broad activation region with a width of 50 to 100 amino acids while the bacterial and oomycete sequences create a some smaller grouped peaks in a broader region which were each around 20 amino acids. We examined further the sequences under the largest peaks, specifically for bacterial sequences we used amino acids 25 to 50, for fungal sequences we used amino acids 35 to 80 and for oomycete sequences we used amino acids 15 to 40. Compositional analysis of the sequence under the peaks showed no apparent primary sequence conservation or motifs as shown in the logo plots in Fig 11 even within a taxon and data set. Enrichment analysis of different amino acid categories showed some statistically significant (*p <* 0.05) changes in proportions of amino acid types relative to the whole set of sequences in the activated regions (Table 3). Bacterial effectors are depleted of hydrophobic amino acids and enriched in polar and basic amino acids. Fungal effectors are enriched in basic amino acids, with no other differences. The oomycete effectors are most interesting in that their activating regions are depleted in hydrophobic and polar amino acids while enriched in neutral and acidic amino acids.

**Table 3:** Enrichment analysis of amino acid types. Proportions of each amino acid type in the activation region of All Sequences: Effector Sequences, with (Probability). Probability by hypergeometric test of the observed number of amino acid type in the region of activation in effector sequences in our compiled sequence set relative to a background of the same region in all sequences in that taxon.

| Taxon | Hydrophobic | Polar | Neutral | Basic | Acidic |
| --- | --- | --- | --- | --- | --- |
| Bacteria | 0.44 : 0.40 (0.00) | 0.26 : 0.28 (0.00) | 0.09 : 0.09 (0.20) | 0.11 : 0.13 (0.00) | 0.09 : 0.09 (0.16) |
| Fungi | 0.39 : 0.38 (0.06) | 0.29 : 0.28 (0.08) | 0.09 : 0.09 (0.29) | 0.13 : 0.14 (0.00) | 0.10 : 0.11 (0.26) |
| Oomycete | 0.44 : 0.43 (0.01) | 0.28 : 0.27 (0.00) | 0.07 : 0.08 (0.00) | 0.10 : 0.10 (0.07) | 0.10 : 0.12 (0.00) |

**Figure 11:**
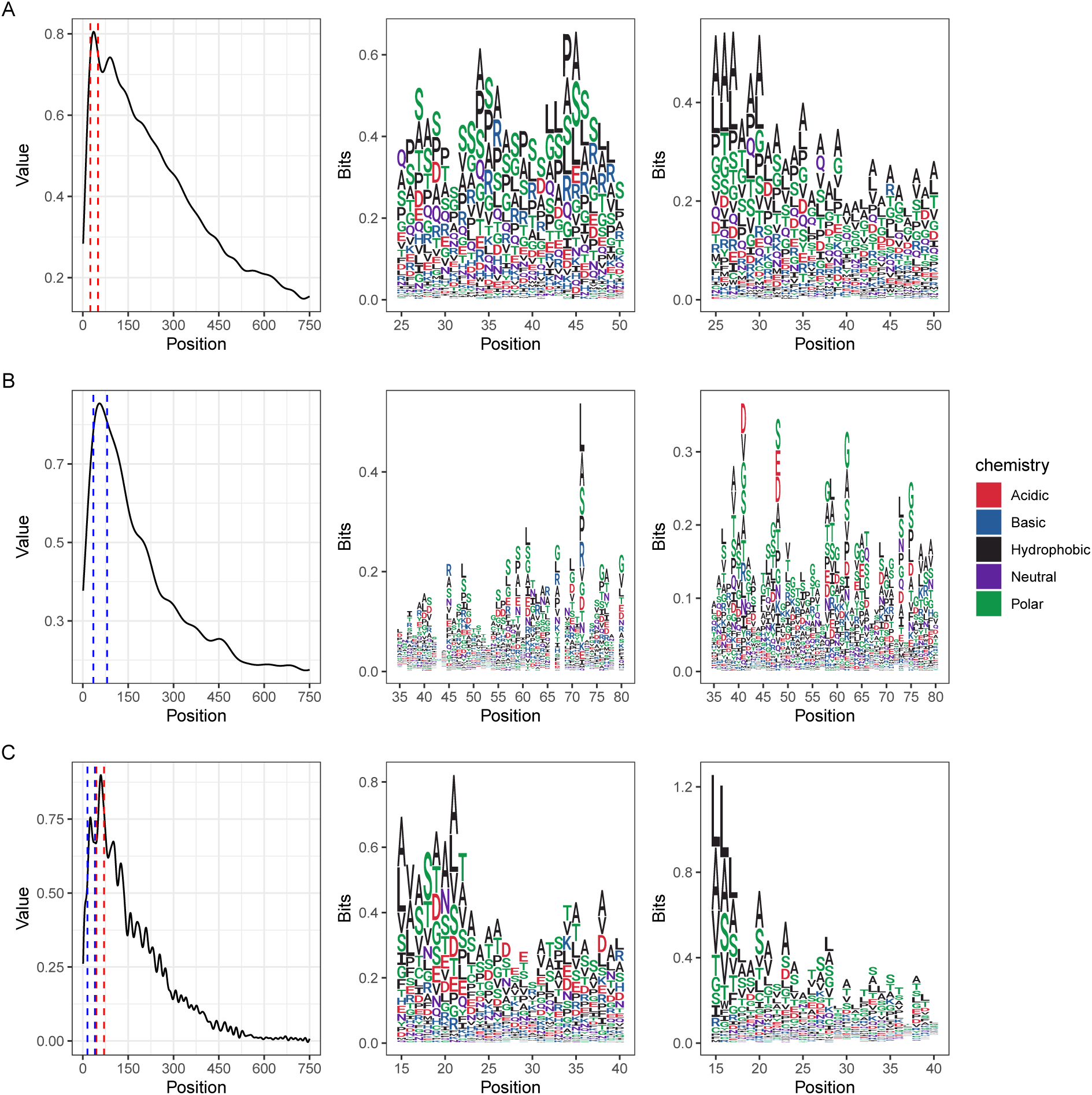
Grad-CAM activation profiles of CNN-LSTM models when run with all datasets; and sequence logo plots of sequence in the activation region within the dotted lines for effector data (left panel) and non-effector data (right panel). A: Models trained on bacterial sequences; B: Models trained on fungal sequences; C: Models trained on oomycete sequences.

### Software Implementation

To make the models useful for developers in their own analytic pipelines we have provided an R package that provides a useful interface to the models. The package and installation instructions are available from GitHub https://ruthkr.github.io/deepredeff

## Discussion

We have compiled a set of known experimentally validated effectors from phytopathogenic bacteria, fungi and oomycetes and used them as positive training examples with which to tune a range of CNN classifiers, the sequences are all taken from the database PHI-Base, a manually curated database that aims to search all the literature on pathogen and host interactions (Urban et al., 2019). The data in PHI-Base is complete as far as 2013 at the moment, thus the phytopathogen effectors we have collected should be all those shown experimentally to have an effect on a host that have been reported on since that time. We chose this set as we believed that this would give us the most reliable and unbiased set of effectors on which we could train learners. That is not to say that the set itself cannot be biased and that the set does not introduce any bias into the classifications of our learners. Sources of bias in our sequence set include the time limits on what has so far been included in PHI-Base, any effectors known but not reported on in this seven year time period cannot be represented in the models. The species of phytopathogens represented in the set also create bias, the effectors are not selected from species sampled proportionally or randomly but instead are those that trends in effector research over the last seven years have brought to focus. In particular, species that have had genome sequencing projects over this time are over-represented. There may also be some echoes of other methods previously applied, effectors studied experimentally must be identified first as hypothetical effectors, usually with the aid of computational tools whose models are themselves biased towards the sort of sequence that we already know. The effectors in the literature may therefore be enriched with respect to known features and results from classifiers should be interpreted with this in mind. We are not easily able to quantify any bias, but the greater generalisability of our models over the others tested gives us reason to believe that the sequence set trained on was broad enough for good models to be developed.

A common misconception of deep learning models is that training datasets need to be extremely large, in practice training data need only to be large with respect to the model size. Here we have coupled small data sets (tens or hundreds of training examples) with small models. The layers and architectures of the models presented here are much smaller than those typically used in large machine vision projects for example. Yet the small models remain useful and have predictive power, indicating a definite role for deep learning approaches in the work of bioinformaticians without truly massive datasets.

Training the models proved to be computationally expensive, the architectures used have a large number of parameters and hyperparameters to be optimised (see Table S4 for a breakdown) and although only a small fraction of the possible hyperparameter and parameter space was explored we compared 3000 models, at a run time of around 144 minutes for CNN-LSTM, 57.2 minutes for CNN-GRU, and 45 minutes for both GRU Embedding and LSTM Embedding. For these relatively small data sets, a significant amount of specialised GPU compute power was required.

The models we created performed exceptionally on the PHI-Base hold-out validation data set of phytopathogenic effector sequences. The greater than 80 % classification accuracy for the fungal and oomycete models is an excellent accuracy on such sequences and our models outperformed the other classifiers tested by large margins on the PHI-Base fungal, no other machine learning method has been reported to have performed as well on phytopathogen sequences. The greater than 97 % accuracy we observed in our model trained on bacterial phytopathogen effectors is also exceptional and similar to what Xue *et al* (Xue et al., 2018) showed in human bacterial pathogen effectors. When we evaluated the proportion of effectors each of our and other classifiers could find in data used to create the other classifiers, we found that our bacterial model outperformed the bacterial models again. A slightly different picture emerged when we compared our method with EffectorP 1 and 2 on fungal data. Both versions of Effector P outperformed our model on training data used to create Effector P in the first instance but identical predictions were made by all fungal classifiers on the validation set provided by Effector P 2. This was coincident with a large drop in accuracy from training to validation data by Effector P 1 and 2. Combined with observations on our PHI-Base data on higher numbers of false positives from Effector P, we conclude that Effector P is over-fitted slightly on its training data and that our model is likely to be more generally accurate. We expect that the tools used in concert will provide very good classification and prediction of fungal effectors. Our model also performed worse on the data used to test the untrained heuristic oomycete RXLR sequence detector EffectR, finding three of six, relative to EffectR’s five of six. This is likely due to the six test sequences being exclusively RXLR and our model being trained on effectors in general, the presence of RXLR not being diagnostic of effectorness in our model means Researchers hoping to find exclusively RXLR containing proteins would be advised to use EffectR, those hoping to find effectors beyond RXLRs may find utility with our oomycete model.

In developing a negative training data set that contained secreted non-effector proteins we hoped to decouple the identification of effector from identification of secreted protein as a proxy for effector. We believe that we have been succesful at this, the models we developed do identify effectors against a background of secreted proteins, indicating that they have some internal representation of what makes an effector different from a secreted peptide. By examining the activation maps of the CNN models, we learned that the maximum activations in the models remains in the N-terminus of the proteins, coincident with the expected positions of secretions signals and is relatively narrow (about 25 amino acids. We also noted that there is no typical primary sequence motif that can be identified, the identity of the amino acids themselves does not seem important. We did find that various categories of amino acid were enriched or depleted significantly in the narrow activation region of effectors, relative to the same area of non-effector secreted proteins. A favouring of functional properties of the amino acids of the effector over the actual identity may be reflective of the many functional types origins of effectors. It may be that the N-terminal activation region in the effectors represents non-canonical secretion-compatible signal in the effector proteins. In order to evolve towards a useful secretion signal from an arbitrary point it may be enough to have a region that satisfies some chemical property that makes it secretable, yet is different enough in effectors to separate them from the secreted proteins by a deep learning algorithm.

Finally, we have made our models available in an R package *deepredeff* - from ‘deep learning prediction of effector’. This can be obtained from GitHub as shown in methods, installation and usage instructions are available in the documentation provided there. The R package allows the user to run the sequences from various sequence format files against a selected model and obtain a score that each sequence is an effector according to the model. Summaries and plots are provided for the user when working on large numbers of sequences. The package integrates well with Bioconductor. For those wishing to use Python to make use of our models, we provide the models as HDF5 files from TensorFlow that can be used in that toolkit.

## Supplementary Tables

**Table S1:** Bacterial species contributing to the bacterial effector sequence set retrieved from PHI-Base

| Species | Sequences Recovered |
| --- | --- |
| *Clavibacter michiganensis* | 2 |
| *Cystobacter fuscus* | 1 |
| *Erwinia amylovora* | 10 |
| *Pantoea stewartii* | 2 |
| *Pseudomonas cichorii* | 1 |
| *Pseudomonas savastanoi* | 1 |
| *Pseudomonas syringae* | 72 |
| *Ralstonia solanacearum* | 46 |
| *Xanthomonas axonopodis* | 4 |
| *Xanthomonas campestris* | 19 |
| *Xanthomonas citri* | 5 |
| *Xanthomonas oryzae* | 24 |
| *Xylella fastidiosa* | 3 |

**Table S5:**
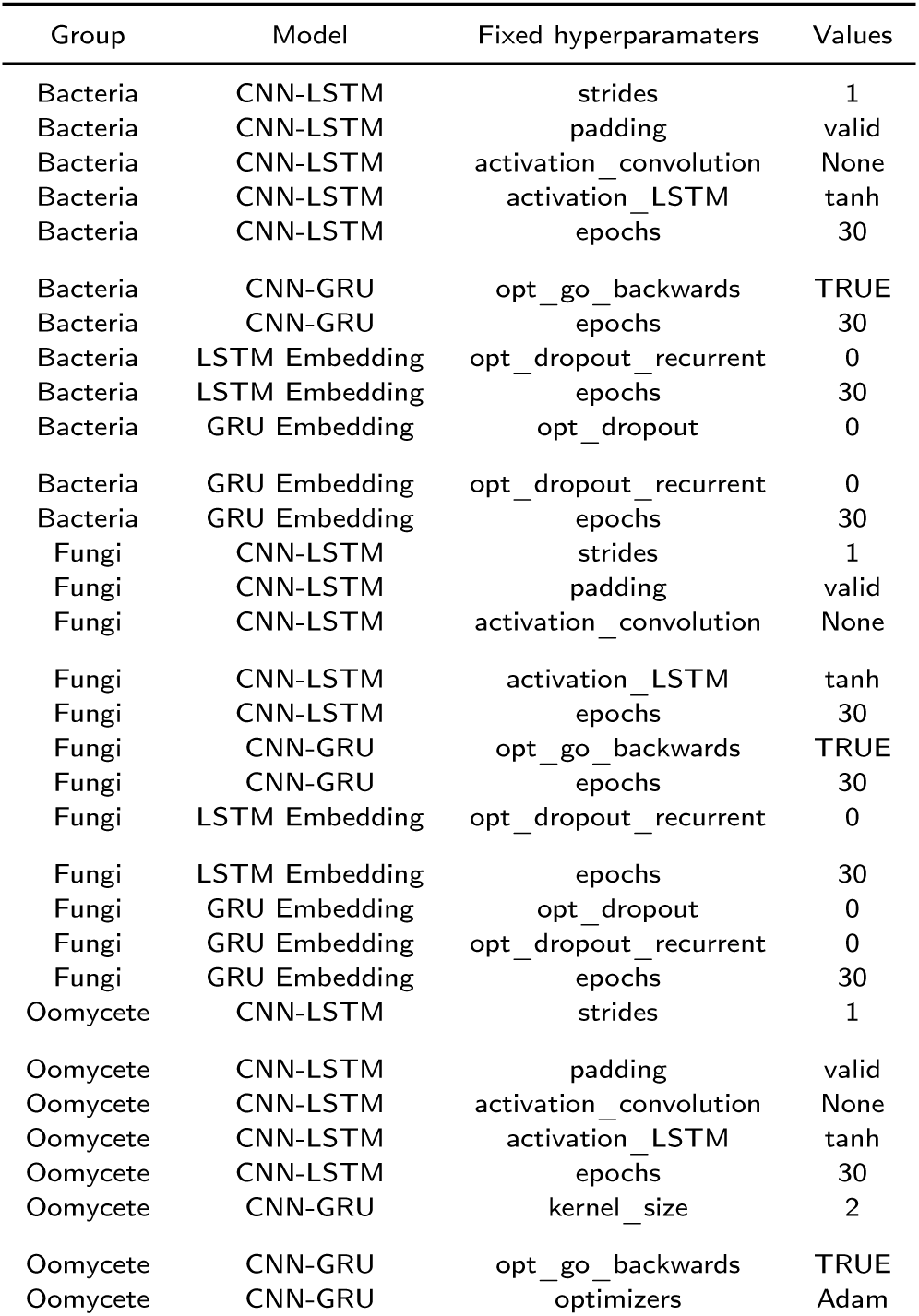
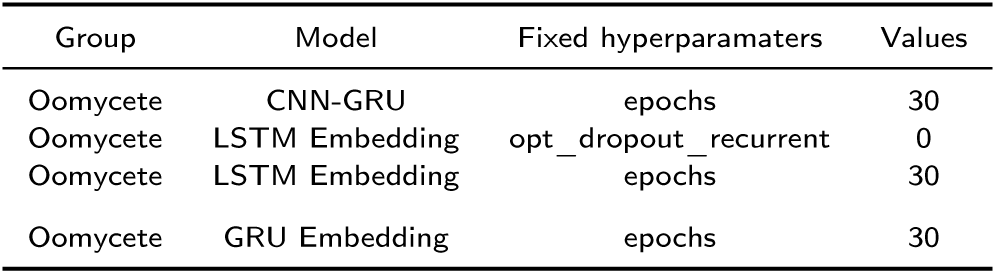
Hyperparameters fixed in automatic scanning of hyperparameter space for each model

**Table S6:** Hyperparameters tuned in manual scanning of hyperparameter space for each model

| Group | Model | Tuned hyperparamaters | Values |
| --- | --- | --- | --- |
| Bacteria | CNN-LSTM | filters | 4, 8, 16 |
| Bacteria | CNN-LSTM | filters_LSTM | 4, 8, 16 |
| Bacteria | CNN-LSTM | optimizers | Adam, Adadelata |
| Bacteria | CNN-LSTM | number_hidden_units | 4, 8 |
| Bacteria | CNN-LSTM | batch_size | 8, 16, 32 |
| Bacteria | CNN-GRU | filter_conv | 8, 16, 32 |
| Bacteria | CNN-GRU | kernel_size | 1, 2 |
| Bacteria | CNN-GRU | maxpool_size | 2, 3 |
| Bacteria | CNN-GRU | activation_conv | None, relu |
| Bacteria | CNN-GRU | gru_hidden_units | 8, 16, 32 |
| Bacteria | CNN-GRU | opt_dropout | 0, 0.25 |
| Bacteria | CNN-GRU | opt_dropout_recurrent | 0, 0.25 |
| Bacteria | CNN-GRU | reg_rate | 0.01, 0.001 |
| Bacteria | CNN-GRU | optimizers | Adam, sgd |
| Bacteria | CNN-GRU | batch_size | 8, 16 |
| Bacteria | LSTM Embedding | outputdim | 16, 32, 64 |
| Bacteria | LSTM Embedding | lstm_hidden_units | 16, 32 |
| Bacteria | LSTM Embedding | opt_dropout | 0, 0.25 |
| Bacteria | LSTM Embedding | opt_go_backwards | TRUE, FALSE |
| Bacteria | LSTM Embedding | reg_rate | 0.01, 0.001 |
| Bacteria | LSTM Embedding | optimizers | Adam, sgd |
| Bacteria | LSTM Embedding | batch_size | 8: 16 |
| Bacteria | GRU Embedding | outputdim | 32, 48 |
| Bacteria | GRU Embedding | gru_hidden_units | 8, 16, 32 |
| Bacteria | GRU Embedding | opt_go_backwards | TRUE, FALSE |
| Bacteria | GRU Embedding | reg_rate | 0.01, 0.001 |
| Bacteria | GRU Embedding | optimizers | 'sgd', 'Adam', 'Adadelata' |
| Bacteria | GRU Embedding | batch_size | 16, 32 |
| Fungi | CNN-LSTM | filters | 4, 8, 16 |
| Fungi | CNN-LSTM | filters_LSTM | 4, 8, 16 |
| Fungi | CNN-LSTM | optimizers | Adam, Adadelata |
| Fungi | CNN-LSTM | number_hidden_units | 4, 8 |
| Fungi | CNN-LSTM | batch_size | 4, 8, 16 |
| Fungi | CNN-GRU | filter_conv | 8, 16 |
| Fungi | CNN-GRU | kernel_size | 1, 2 |
| Fungi | CNN-GRU | maxpool_size | 2, 3 |
| Fungi | CNN-GRU | activation_conv | None, relu |
| Fungi | CNN-GRU | gru_hidden_units | 8, 16 |
| Fungi | CNN-GRU | opt_dropout | 0, 0.25 |
| Fungi | CNN-GRU | opt_dropout_recurrent | 0, 0.25 |
| Fungi | CNN-GRU | reg_rate | 0.01, 0.001 |
| Fungi | CNN-GRU | optimizers | Adam, sgd |
| Fungi | CNN-GRU | batch_size | 4, 8 |
| Fungi | LSTM Embedding | outputdim | 16, 32, 64 |
| Fungi | LSTM Embedding | lstm_hidden_units | 16, 32 |
| Fungi | LSTM Embedding | opt_dropout | 0, 0.25 |
| Fungi | LSTM Embedding | opt_go_backwards | TRUE, FALSE |
| Fungi | LSTM Embedding | reg_rate | 0.01, 0.001 |
| Fungi | LSTM Embedding | optimizers | Adam, sgd |
| Fungi | LSTM Embedding | batch_size | 4, 8 |
| Fungi | GRU Embedding | outputdim | 8, 16, 32 |
| Fungi | GRU Embedding | gru_hidden_units | 8, 16, 32 |
| Fungi | GRU Embedding | opt_go_backwards | TRUE, FALSE |

| Group | Model | Tuned hyperpramaters | Values |
| --- | --- | --- | --- |
| Fungi | GRU Embedding | reg_rate | 0.01, 0.001 |
| Fungi | GRU Embedding | optimizers | 'sgd', 'Adam', 'Adadelata' |
| Fungi | GRU Embedding | batch_size | 4, 8 |
| Oomycete | CNN-LSTM | filters | 4, 8, 16 |
| Oomycete | CNN-LSTM | filters_LSTM | 4, 8, 16 |
| Oomycete | CNN-LSTM | optimizers | Adam, Adadelata |
| Oomycete | CNN-LSTM | number_hidden_units | 4, 8 |
| Oomycete | CNN-LSTM | batch_size | 4, 8, 16 |
| Oomycete | CNN-GRU | filter_conv | 8, 16, 32 |
| Oomycete | CNN-GRU | maxpool_size | 2, 3 |
| Oomycete | CNN-GRU | activation_conv | None, relu |
| Oomycete | CNN-GRU | gru_hidden_units | 8, 16, 32 |
| Oomycete | CNN-GRU | opt_dropout | 0, 0.25 |
| Oomycete | CNN-GRU | opt_dropout_recurrent | 0, 0.25 |
| Oomycete | CNN-GRU | reg_rate | 0.01, 0.001 |
| Oomycete | CNN-GRU | batch_size | 4, 8 |
| Oomycete | LSTM Embedding | outputdim | 16, 32, 64 |
| Oomycete | LSTM Embedding | lstm_hidden_units | 16, 32 |
| Oomycete | LSTM Embedding | opt_dropout | 0, 0.25 |
| Oomycete | LSTM Embedding | opt_go_backwards | TRUE, FALSE |
| Oomycete | LSTM Embedding | reg_rate | 0.01, 0.001 |
| Oomycete | LSTM Embedding | optimizers | Adam, sgd |
| Oomycete | LSTM Embedding | batch_size | 4, 8 |
| Oomycete | GRU Embedding | outputdim | 32, 48, 16 |
| Oomycete | GRU Embedding | gru_hidden_units | 8, 16, 32, 64 |
| Oomycete | GRU Embedding | opt_dropout | 0, 0.25, 0.5 |
| Oomycete | GRU Embedding | opt_dropout_recurrent | 0, 0.25, 0.5 |
| Oomycete | GRU Embedding | opt_go_backwards | TRUE, FALSE |
| Oomycete | GRU Embedding | reg_rate | 0.01, 0.001 |
| Oomycete | GRU Embedding | optimizers | 'sgd', 'Adam', 'Adadelata' |
| Oomycete | GRU Embedding | batch_size | 4, 8 |

**Table S7:**
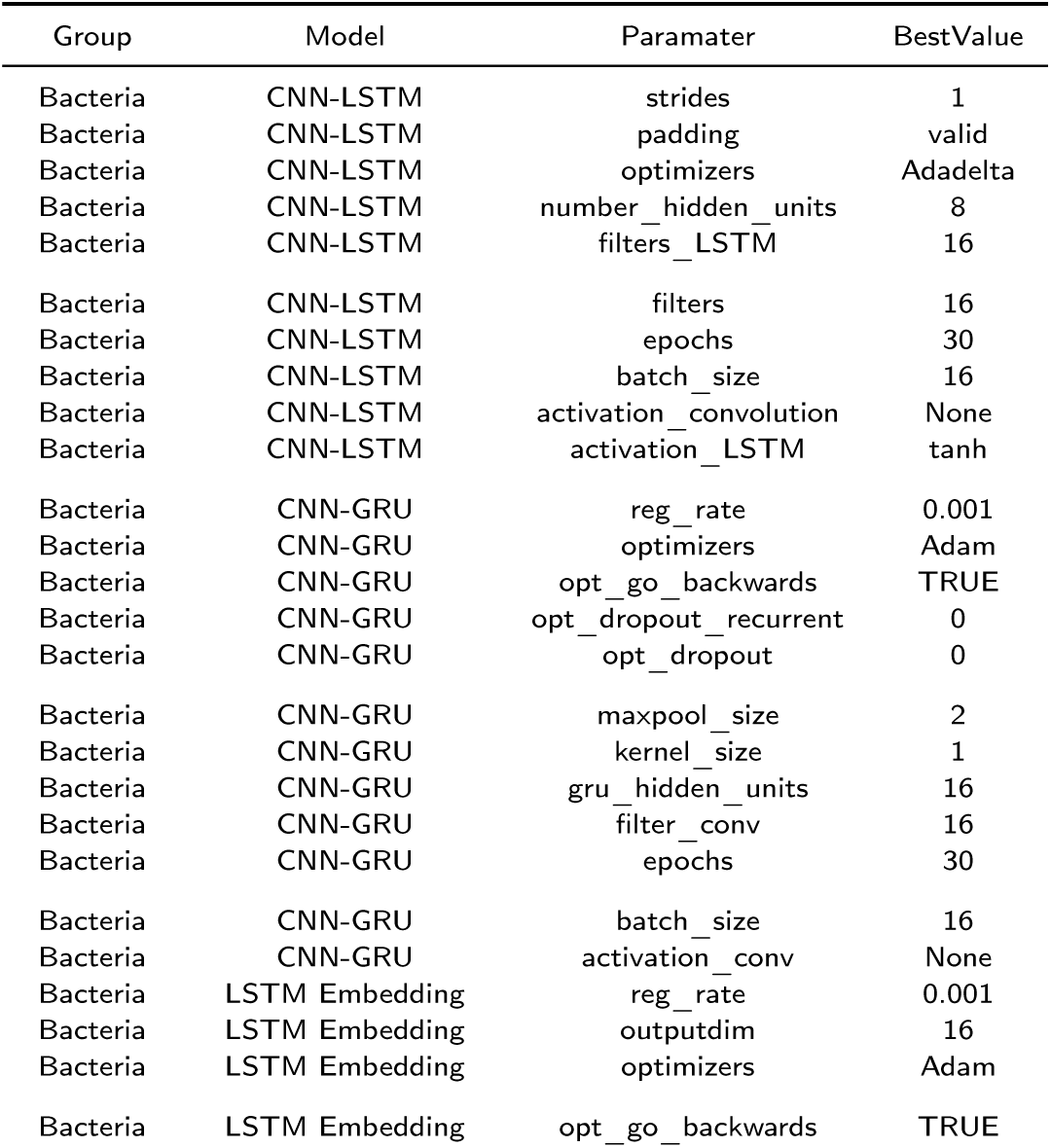
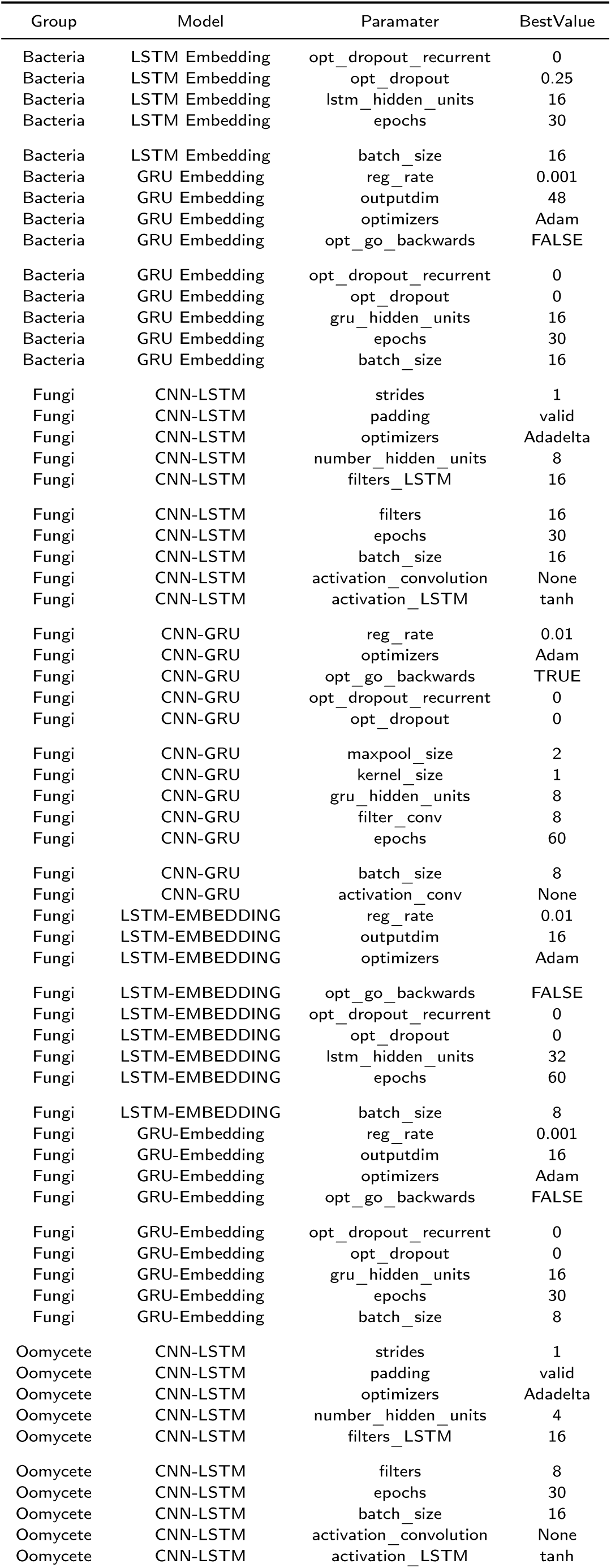
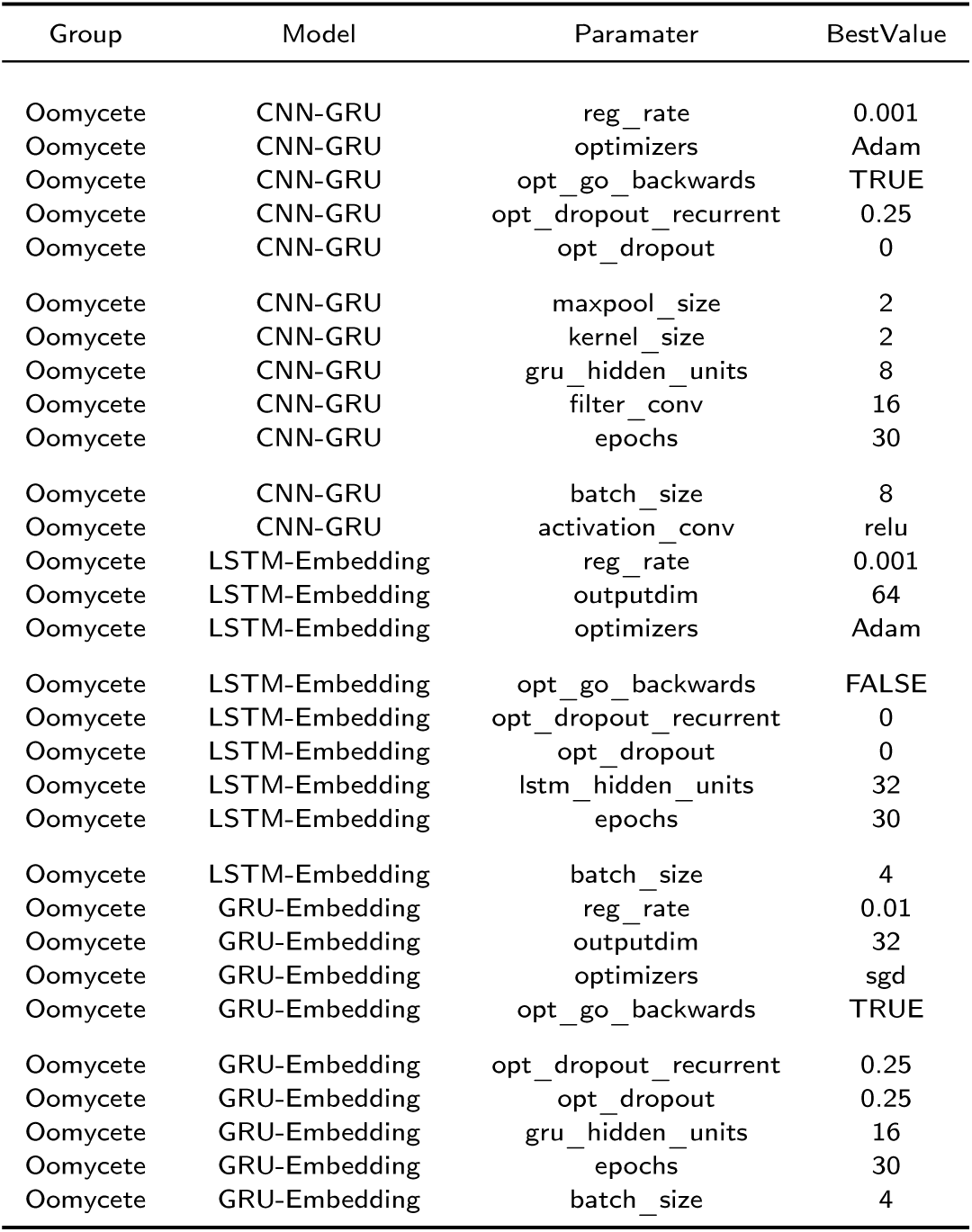
Hyperparameters used in automatic scanning of hyperparameter space and best performing parameters values for each model

| Group | Model | Paramater | BestValue |
| --- | --- | --- | --- |
| Bacteria | CNN-LSTM | strides | 1 |
| Bacteria | CNN-LSTM | padding | valid |
| Bacteria | CNN-LSTM | optimizers | Adadelata |
| Bacteria | CNN-LSTM | number_hidden_units | 8 |
| Bacteria | CNN-LSTM | filters_LSTM | 16 |
| Bacteria | CNN-LSTM | filters | 16 |
| Bacteria | CNN-LSTM | epochs | 30 |
| Bacteria | CNN-LSTM | batch_size | 16 |
| Bacteria | CNN-LSTM | activation_convolution | None |
| Bacteria | CNN-LSTM | activation_LSTM | tanh |
| Bacteria | CNN-GRU | reg_rate | 0.001 |
| Bacteria | CNN-GRU | optimizers | Adam |
| Bacteria | CNN-GRU | opt_go_backwards | TRUE |
| Bacteria | CNN-GRU | opt_dropout_recurrent | 0 |
| Bacteria | CNN-GRU | opt_dropout | 0 |
| Bacteria | CNN-GRU | maxpool_size | 2 |
| Bacteria | CNN-GRU | kernel_size | 1 |
| Bacteria | CNN-GRU | gru_hidden_units | 16 |
| Bacteria | CNN-GRU | filter_conv | 16 |
| Bacteria | CNN-GRU | epochs | 30 |
| Bacteria | CNN-GRU | batch_size | 16 |
| Bacteria | CNN-GRU | activation_conv | None |
| Bacteria | LSTM Embedding | reg_rate | 0.001 |
| Bacteria | LSTM Embedding | outputdim | 16 |
| Bacteria | LSTM Embedding | optimizers | Adam |
| Bacteria | LSTM Embedding | opt_go_backwards | TRUE |
| Bacteria | LSTM Embedding | opt_dropout_recurrent | 0 |
| Bacteria | LSTM Embedding | opt_dropout | 0.25 |
| Bacteria | LSTM Embedding | lstm_hidden_units | 16 |
| Bacteria | LSTM Embedding | epochs | 30 |
| Bacteria | LSTM Embedding | batch_size | 16 |
| Bacteria | GRU Embedding | reg_rate | 0.001 |
| Bacteria | GRU Embedding | outputdim | 48 |
| Bacteria | GRU Embedding | optimizers | Adam |
| Bacteria | GRU Embedding | opt_go_backwards | FALSE |
| Bacteria | GRU Embedding | opt_dropout_recurrent | 0 |
| Bacteria | GRU Embedding | opt_dropout | 0 |
| Bacteria | GRU Embedding | gru_hidden_units | 16 |
| Bacteria | GRU Embedding | epochs | 30 |
| Bacteria | GRU Embedding | batch_size | 16 |
| Fungi | CNN-LSTM | strides | 1 |
| Fungi | CNN-LSTM | padding | valid |
| Fungi | CNN-LSTM | optimizers | Adadelta |
| Fungi | CNN-LSTM | number_hidden_units | 8 |
| Fungi | CNN-LSTM | filters_LSTM | 16 |
| Fungi | CNN-LSTM | filters | 16 |
| Fungi | CNN-LSTM | epochs | 30 |
| Fungi | CNN-LSTM | batch_size | 16 |
| Fungi | CNN-LSTM | activation_convolution | None |
| Fungi | CNN-LSTM | activation_LSTM | tanh |
| Fungi | CNN-GRU | reg_rate | 0.01 |
| Fungi | CNN-GRU | optimizers | Adam |
| Fungi | CNN-GRU | opt_go_backwards | TRUE |
| Fungi | CNN-GRU | opt_dropout_recurrent | 0 |
| Fungi | CNN-GRU | opt_dropout | 0 |
| Fungi | CNN-GRU | maxpool_size | 2 |
| Fungi | CNN-GRU | kernel_size | 1 |
| Fungi | CNN-GRU | gru_hidden_units | 8 |
| Fungi | CNN-GRU | filter_conv | 8 |
| Fungi | CNN-GRU | epochs | 60 |
| Fungi | CNN-GRU | batch_size | 8 |
| Fungi | CNN-GRU | activation_conv | None |
| Fungi | LSTM-EMBEDDING | reg_rate | 0.01 |
| Fungi | LSTM-EMBEDDING | outputdim | 16 |
| Fungi | LSTM-EMBEDDING | optimizers | Adam |
| Fungi | LSTM-EMBEDDING | opt_go_backwards | FALSE |
| Fungi | LSTM-EMBEDDING | opt_dropout_recurrent | 0 |
| Fungi | LSTM-EMBEDDING | opt_dropout | 0 |
| Fungi | LSTM-EMBEDDING | lstm_hidden_units | 32 |
| Fungi | LSTM-EMBEDDING | epochs | 60 |
| Fungi | LSTM-EMBEDDING | batch_size | 8 |
| Fungi | GRU-Embedding | reg_rate | 0.001 |
| Fungi | GRU-Embedding | outputdim | 16 |
| Fungi | GRU-Embedding | optimizers | Adam |
| Fungi | GRU-Embedding | opt_go_backwards | FALSE |
| Fungi | GRU-Embedding | opt_dropout_recurrent | 0 |
| Fungi | GRU-Embedding | opt_dropout | 0 |
| Fungi | GRU-Embedding | gru_hidden_units | 16 |
| Fungi | GRU-Embedding | epochs | 30 |
| Fungi | GRU-Embedding | batch_size | 8 |
| Oomycete | CNN-LSTM | strides | 1 |
| Oomycete | CNN-LSTM | padding | valid |
| Oomycete | CNN-LSTM | optimizers | Adadelta |
| Oomycete | CNN-LSTM | number_hidden_units | 4 |
| Oomycete | CNN-LSTM | filters_LSTM | 16 |
| Oomycete | CNN-LSTM | filters | 8 |
| Oomycete | CNN-LSTM | epochs | 30 |
| Oomycete | CNN-LSTM | batch_size | 16 |
| Oomycete | CNN-LSTM | activation_convolution | None |
| Oomycete | CNN-LSTM | activation_LSTM | tanh |

| Group | Model | Parameter | BestValue |
| --- | --- | --- | --- |
| Oomycete | CNN-GRU | reg_rate | 0.001 |
| Oomycete | CNN-GRU | optimizers | Adam |
| Oomycete | CNN-GRU | opt_go_backwards | TRUE |
| Oomycete | CNN-GRU | opt_dropout_recurrent | 0.25 |
| Oomycete | CNN-GRU | opt_dropout | 0 |
| Oomycete | CNN-GRU | maxpool_size | 2 |
| Oomycete | CNN-GRU | kernel_size | 2 |
| Oomycete | CNN-GRU | gru_hidden_units | 8 |
| Oomycete | CNN-GRU | filter_conv | 16 |
| Oomycete | CNN-GRU | epochs | 30 |
| Oomycete | CNN-GRU | batch_size | 8 |
| Oomycete | CNN-GRU | activation_conv | relu |
| Oomycete | LSTM-Embedding | reg_rate | 0.001 |
| Oomycete | LSTM-Embedding | outputdim | 64 |
| Oomycete | LSTM-Embedding | optimizers | Adam |
| Oomycete | LSTM-Embedding | opt_go_backwards | FALSE |
| Oomycete | LSTM-Embedding | opt_dropout_recurrent | 0 |
| Oomycete | LSTM-Embedding | opt_dropout | 0 |
| Oomycete | LSTM-Embedding | lstm_hidden_units | 32 |
| Oomycete | LSTM-Embedding | epochs | 30 |
| Oomycete | LSTM-Embedding | batch_size | 4 |
| Oomycete | GRU-Embedding | reg_rate | 0.01 |
| Oomycete | GRU-Embedding | outputdim | 32 |
| Oomycete | GRU-Embedding | optimizers | sgd |
| Oomycete | GRU-Embedding | opt_go_backwards | TRUE |
| Oomycete | GRU-Embedding | opt_dropout_recurrent | 0.25 |
| Oomycete | GRU-Embedding | opt_dropout | 0.25 |
| Oomycete | GRU-Embedding | gru_hidden_units | 16 |
| Oomycete | GRU-Embedding | epochs | 30 |
| Oomycete | GRU-Embedding | batch_size | 4 |

**Table S2:** Fungal species contributing to the fungal effector sequence set retrieved from PHI-Base

| Species | Sequences Recovered |
| --- | --- |
| *Blumeria graminis* | 14 |
| *Botrytis cinerea* | 1 |
| *Cercospora beticola* | 1 |
| *Colletotrichum orbiculare* | 1 |
| *Dothistroma septosporum* | 1 |
| *Fusarium graminearum* | 1 |
| *Fusarium oxysporum* | 9 |
| *Leptosphaeria maculans* | 6 |
| *Magnaporthe oryzae* | 19 |
| *Penicillium expansum* | 4 |
| *Puccinia striiformis* | 2 |
| *Pyrenophora tritici-repentis* | 2 |
| *Rhynchosporium commune* | 3 |
| *Ustilago maydis* | 25 |
| *Verticillium dahliae* | 6 |
| *Zymoseptoria tritici* | 2 |

**Table S3:** Oomycete species contributing to the oomycete effector sequence set retrieved from PHI-Base

| Species | Sequences Recovered |
| --- | --- |
| *Hyaloperonospora arabidopsidis* | 42 |
| *Phytophthora cactorum* | 1 |
| *Phytophthora infestans* | 20 |
| *Phytophthora parasitica* | 2 |
| *Phytophthora sojae* | 19 |
| *Pythium aphanidermatum* | 1 |

**Table S4:** List of possible combination of hyperparamaters setting for each model

| Group | Model | Possible combination |
| --- | --- | --- |
| Bacteria | CNN-LSTM | 108 |
| Bacteria | CNN-GRU | 2,304 |
| Bacteria | LSTM Embedding | 192 |
| Bacteria | GRU Embedding | 144 |
| Fungi | CNN-LSTM | 108 |
| Fungi | CNN-GRU | 1,024 |
| Fungi | LSTM Embedding | 192 |
| Fungi | GRU Embedding | 216 |
| Oomycete | CNN-LSTM | 108 |
| Oomycete | CNN-GRU | 576 |
| Oomycete | LSTM Embedding | 128 |
| Oomycete | GRU Embedding | 2,592 |

